# Evidence for intrinsic DNA dynamics and deformability in damage sensing by the Rad4/XPC nucleotide excision repair complex

**DOI:** 10.1101/2024.04.02.587835

**Authors:** Saroj Baral, Sagnik Chakraborty, Peter J. Steinbach, Debamita Paul, Jung-Hyun Min, Anjum Ansari

## Abstract

Altered DNA dynamics at lesion sites are implicated in how DNA repair proteins sense damage within genomic DNA. We examined DNA dynamics related to damage recognition by Rad4 (yeast ortholog of XPC), which recognizes diverse lesions from environmental mutagens and initiates nucleotide excision repair. Using laser temperature-jump (T-jump), we measured the dynamics of DNA containing 3 base-pair mismatches recognized specifically by Rad4 *in vitro*. The T-jump kinetics traces measured using a cytosine-analog FRET pair, together with rigorous comparison with equilibrium measurements, enabled conformational dynamics to be revealed beyond the T-jump observation window of ∼20 µs – 50 ms. AT-rich nonspecific sites (matched or mismatched) exhibited dynamics primarily within the T-jump window, albeit with some amplitude in “missing” fast (< 20 µs) kinetics. The fast-kinetics amplitudes increased dramatically for specific sites, which were further distinguished by additional ‘missing’ amplitude in slow (> 50 ms) kinetics at elevated temperatures. We posit that the rapid (µs-ms) fluctuations help stall a diffusing protein at AT-rich/damaged sites and that the >50-ms kinetics reflect a propensity for specific DNA to adopt unwound/bent conformations that may resemble Rad4-bound structures. These studies provide compelling evidence for unusual DNA dynamics and deformability that likely govern how Rad4 senses DNA damage.

## INTRODUCTION

How DNA damage sensing proteins search for and identify damaged sites within undamaged genomic DNA remains unresolved. While many studies have focused on how these proteins juggle three-dimensional (3D) and one-dimensional (1D) diffusion to optimize the search process, the mechanism by which these proteins recognize damage when in the vicinity is not well understood. In particular, the role DNA plays in recruiting proteins to potential damage sites remains poorly characterized. Altered DNA dynamics at lesion sites are implicated in damage sensing, but characterizing these dynamics has been a challenge. We examined DNA dynamics in the context of damage recognition by Rad4 (yeast ortholog of XPC), which recognizes diverse lesions from UV-damage or other genotoxins and initiates nucleotide excision repair (NER). Previous structural studies showed that when Rad4 binds to a lesion site, it unwinds the DNA at that site, flips out two nucleotide pairs that include the lesion, and inserts a β-hairpin into the gap in the DNA to stabilize this so-called ‘open’ conformation (1,2). Notably, in this recognition complex, Rad4 does not make direct contact with the damaged nucleotides but interacts exclusively with the nucleotides on the opposite strand (Figure 1). This ‘indirect readout’ enables the protein to bind to a variety of structurally dissimilar lesions that are repaired by NER. Rad4 also binds *in vitro* to DNA sites with 2 or 3 base pair (bp) mismatches with affinities similar to *bona fide* NER lesions and forms the same ‘open’ complex as with NER lesions, making these mismatched sites suitable model systems for exploring the damage sensing mechanisms (3–5).

**Figure 1.**
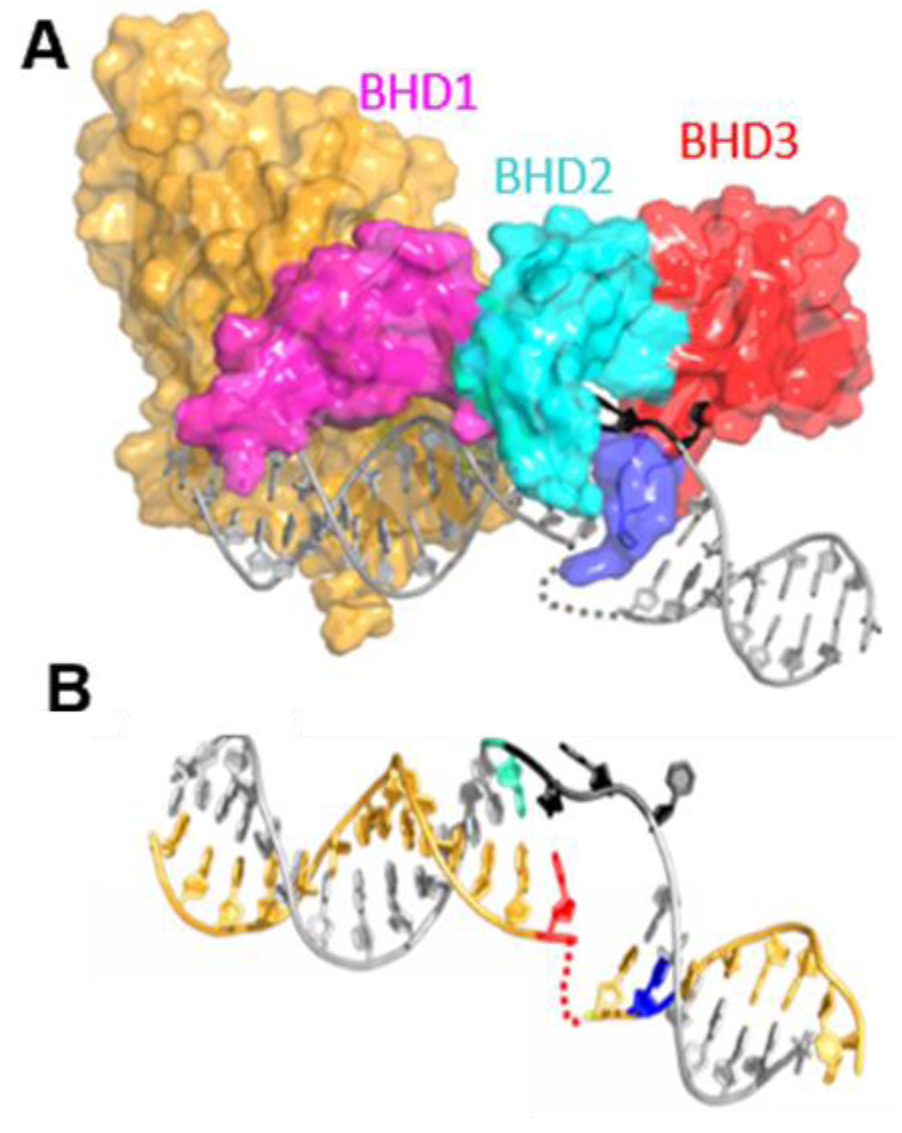
NER damage sensing protein Rad4 (yeast ortholog of XPC. (A) Structure of Rad4-bound specific complex (PDB ID: 2QSH). The β-hairpin-domains (BHD) are indicated; β-hairpin from BHD2 (blue) is inserted into the DNA at the lesion site; the damaged nucleotides are flipped out and disordered in the crystal (gray dotted line); undamaged nucleotides on the opposite strand are flipped towards the protein (black). (B) Structure of DNA in the Rad4-bound specific complex. The positions of 3 bp mismatched nucleotides are indicated in red and black. The positions of the donor (tC°, blue) and acceptor (tC_nitro_, cyan), incorporated in the DNA for the fluorescence measurements presented in this study, are also shown.

Remarkably, Rad4 when covalently tethered to undamaged DNA is also able to flip out nucleotides in a manner akin to the specific complex (3), although certain sequences are more prone to being ‘opened’ than others (6). The observation that Rad4 could flip out nucleotides from within undamaged sites, if held at that site long enough, led to the proposal of a ‘kinetic gating’ mechanism for damage recognition by Rad4/XPC. In this mechanism, the ability of a freely diffusing protein to flip nucleotides at any given site is governed by the ratio of the probability that Rad4 will flip nucleotides when sitting at that site versus the probability that it will diffuse over to an adjacent site. This mechanism allows Rad4, while searching for damage, to more efficiently flip nucleotides at lesion sites compared with undamaged sites, since the rates for nucleotide flipping are expected to be much faster at helix-destabilizing lesions compared with stable, undamaged sites.

In a previous study, we employed laser temperature-jump (T-jump) spectroscopy to unveil Rad4 induced DNA unwinding and nucleotide-flipping kinetics on DNA constructs with 2- or 3-bp mismatches (3,4). DNA conformational dynamics in response to a T-jump perturbation were monitored using either 2-aminopurine (2AP; fluorescent analog of adenine) placed within the mismatch, to probe directly nucleotide flipping, or a cytosine analog FRET pair (tC° and tC_nitro_) placed on either side of the mismatched site, to probe DNA unwinding. The T-jump measurements were carried out on DNA with TTT/TTT mismatch that is specifically recognized by Rad4 and TAT/TAT mismatch that is not recognized by Rad4 and hence served as a nonspecific control together with the matched counterparts. These measurements indicated a ‘twist-open’ mechanism for mismatch recognition by Rad4, with nonspecific unwinding on ∼100–500 µs timescale attributed to interrogation dynamics followed by nucleotide flipping on timescales of ∼5–10 ms to form the recognition complex (4). Nucleotide flipping times from within undamaged DNA are predicted to be several orders of magnitude slower (3), although they have not been directly measured.

These earlier T-jump studies focused on readily observed Rad4-facilitated DNA conformational dynamics and did not report on intrinsic DNA dynamics in the absence of Rad4, either because DNA-only dynamics fell outside of the T-jump observation window or the fluorescent probes, depending on where they were placed on the DNA, were insensitive to smaller amplitude DNA-only dynamics. In a follow-up study we employed fluorescence lifetime measurements with the tC° and tC_nitro_ FRET probes to examine the equilibrium distributions of thermally accessible DNA conformations and uncovered evidence for intrinsic fluctuations in mismatched DNA specifically recognized by Rad4 (5). This study, which also included a third construct with a CCC/CCC mismatch, showed that the specific constructs (TTT/TTT and CCC/CCC) exhibited a broader range of accessible DNA conformations than their matched counterparts even in the absence of Rad4, while the nonspecific TAT/TAT construct remained predominantly B-DNA-like. Indeed, the CCC/CCC construct, which had the highest affinity for Rad4 among all the mismatches studied, exhibited the broadest conformational heterogeneity, with DNA conformations ranging from B-DNA to highly distorted conformations that resembled those seen in crystal structures of Rad4-DNA specific complexes. Upon Rad4 binding, the conformational distributions of the specific constructs shifted further towards the more distorted conformations. While these studies uncovered that mismatched DNA specifically recognized by Rad4 could adopt distorted conformations from spontaneous thermal fluctuations, they did not reveal the timescales of these fluctuations, since lifetime studies provide only a snapshot of DNA conformations.

We therefore returned to laser T-jump for a renewed examination of the dynamics of spontaneous DNA fluctuations on all three mismatched constructs and their matched counterparts. Elucidating the amplitudes and timescales of intrinsic DNA fluctuations is important for piecing together the puzzle of how damage sensing proteins that are known to diffuse relatively rapidly on DNA during the search process slow down sufficiently to engage with potential damage. Single molecule imaging studies of 1D diffusion of Rad4 and XPC on undamaged and damaged DNA have revealed multiple modes of motion for the protein, ranging from free diffusion to constrained diffusion to being immobilized, at least to within the spatial resolution of these measurements (7,8). Notably, the sites of constrained diffusion or apparently stalled proteins were found to correlate with AT-rich sites on DNA, implicating enhanced breathing/opening of the base pairs at AT-rich versus GC-rich sites as potential mechanism for inducing a conformational change in the protein and thereby altering its diffusion mode. We find that DNA with even a 3 bp stretch of As and Ts, whether matched or mismatched, exhibits conformational relaxation dynamics in response to a T-jump perturbation, albeit with larger amplitudes for mismatched versus matched. These dynamics were observed over a broad range of timescales within the T-jump observation window of ∼20 µs to ∼50 ms. Additionally, using a new analysis approach with quantitative comparison between the T-jump observed conformational relaxation amplitudes with those predicted from equilibrium thermal profiles, we uncovered evidence for ‘missing’ (or ‘hidden’) kinetics that fell outside the T-jump-resolved observation window. We uncovered ‘missing’ fast (< 20 µs) kinetics, the amplitudes of which were significantly larger for mismatched sites specifically recognized by Rad4. Furthermore, for the specific constructs, we also uncovered ‘missing’ slow (> 50 ms) kinetics; matched or mismatched-but-nonspecific sites did not exhibit any missing slow amplitudes. Matched DNA containing a stretch of Gs and Cs was not readily perturbed by T-jump and yielded very small amplitude kinetics, if any. Taken together, we propose that the micro-to-millisecond dynamics observed in all AT-rich (matched or mismatched) and all specific constructs are the ones responsible for engaging the protein and interrupting or altering its diffusing mode, while the slow (> 50 ms) kinetics suggest a propensity for specific DNA to spontaneously access the more severely distorted/unwound conformations – perhaps with nucleotides partially flipped out – albeit with high free energy barriers. Rad4, once engaged, lowers the barrier for full distortion of the DNA and forms the recognition complex within ∼25–50 ms. These studies provide the first in-depth view of the range of intrinsic DNA dynamics that Rad4 likely senses during interrogation and unveils features of the dynamics that distinguish specific from nonspecific sites.

## MATERIAL AND METHODS

All details of sample preparation, data acquisition, and data analysis are described in the Supplementary Data. Here we briefly describe the materials and methods.

### DNA and protein samples

tC°-tC_nitro_ –labeled DNA samples (see Supplementary Figure S1) were purchased from TriLink Biotechnologies and prepared as described in SI Methods 1.1. Rad4 samples were prepared as described in SI Methods 1.2

### Equilibrium measurements

The steady-state fluorescence emission spectra for donor-only (DNA_D) and donor-acceptor-labeled (DNA_DA) samples were measured on a FluoroMax4 spectrofluorimeter (Jobin Yvon, Inc., NJ); FRET efficiencies (FRET E) were obtained from the measured spectra as described in SI Methods 1.3. Corresponding fluorescence decay curves were measured with a PicoMaster fluorescence lifetime instrument (HORIBA-PTI, London, Ontario, Canada); fluorescence lifetime distributions and FRET efficiencies were obtained from the measured decay traces as described in SI Methods 1.4.

### Laser T-jump measurements

tC°-tC_nitro_ –labeled DNA samples were perturbed by inducing a rapid 5–10 ℃ T-jump using 10 ns IR laser pulses at 1550 nm. The details of the T-jump spectrometer are as described previously (4). Salient description of the spectrometer, data acquisition and analysis is in SI Methods 1.5-1.8. Measurements were performed with the initial temperature set in the temperature range 10 − 35 ℃, with the final temperatures, after the T-jump, spanning 14 − 41℃. The fluorescence of the FRET donor (tC°) as detected in the presence of the acceptor (tC_nitro_) was used as the reporter of the conformational relaxation kinetics in response to the T-jump perturbation. Measurements were performed on donor-only samples (denoted as DNA_D) and donor-acceptor-labeled samples (denoted as DNA_DA), with and without Rad4. The maximum entropy method (MEM), described in SI Methods 1.9, was used to infer the distribution of rate constants from the relaxation traces (9,10). The amplitudes of the observed relaxation dynamics were compared with the amplitudes expected from steady-state measurements to estimate the extent of the relaxation kinetics not resolved within the T-jump window – the so-called ‘missing’ amplitudes – as described in SI Methods 1.10.

## RESULTS

### AT-rich nonspecific sites and mismatched specific sites are readily perturbed by a temperature increase

The alterations in DNA conformations with increase in temperature were examined using equilibrium FRET measurements on tC°-tC_nitro_–labeled DNA constructs, with and without Rad4. The DNA sequence and placement of the labels are summarized in Supplementary Figure S1. The equilibrium FRET measurements and the T-jump studies were performed in the temperature range 10–40 ℃. Previous studies demonstrated that Rad4 bound to DNA remains stable up to 40 ℃ (3,4). FRET measurements were performed both with steady-state fluorescence and fluorescence lifetime. Representative donor emission spectra measured under steady-state conditions are shown in Supplementary Figure S2. The dramatic suppression of the donor emission intensities in the presence of the acceptor (DNA_DA) compared with donor-only emission intensities (DNA_D) is the result of FRET efficiency (E) between the probes. Corresponding measurements from fluorescence lifetime measurements are shown in Supplementary Figure S3. The FRET E versus temperature profiles for each of the constructs measured by the two different approaches yielded consistent results (SI Methods 1.3–1.4 and Supplementary Figure S1).

Below, we briefly review the salient results from our earlier studies on these DNA constructs that set the stage for interpreting the T-jump measurements reported here (5). First, the FRET E values for matched DNA constructs were in the range of ∼0.92 − 0.94 at 10 ℃, consistent with the FRET of 0.94 computed for a canonical B-DNA structure based on the known distance and relative orientation of cytosines at the position of our probes (5,11). Second, the FRET E values for matched GGG/CCC remained essentially unchanged as the temperature was raised, while the other two matched constructs (AAA/TTT and ATA/TAT) exhibited a slight decrease in FRET with increase in temperature, indicative of more deformable DNA than the GGG/CCC construct (Supplementary Figure S1).

The table in Supplementary Figure S1 summarizes the binding affinities of each of the DNA constructs in this study and their thermal stability. Mismatched CCC/CCC exhibited the highest specificity for Rad4 among the three mismatched constructs; correspondingly, it also exhibited the largest alteration in its average DNA conformation compared with its matched counterpart, as indicated by a significantly different FRET E value compared with GGG/CCC; the conformations were further distorted upon Rad4 binding (Supplementary Figure S1). The TTT/TTT mismatch exhibited lower specificity for Rad4 and was only slightly altered in its average conformation in comparison with AAA/TTT. Rad4 binding further distorted TTT/TTT (lower FRET E) but not as much as CCC/CCC. Finally, like all other mismatched constructs, the nonspecific TAT/TAT exhibited FRET E values that were lower in comparison with the matched ATA/TAT, indicating a mismatch-induced alteration in its DNA conformation. However, Rad4 binding did not alter the mismatched conformation any further, consistent with nonspecific binding to TAT/TAT.

While the average FRET E values are useful as a first look at DNA conformational changes in the presence of mismatches or upon Rad4 binding, the full extent of the DNA conformations are better visualized in the fluorescence lifetime distribution plots (Figure 2). The matched GGG/CCC construct shows a predominantly single-peaked distribution with little or no change upon increase in temperature (Figure 2A), while the mismatched CCC/CCC construct and its Rad4-bound complex exhibit multiple peaks in the lifetime distribution, that shift to longer lifetimes (more distorted conformations) with increasing temperature (Figure 2B-C). Corresponding lifetime distributions for the matched AAA/TTT also show predominantly the B-DNA-like peak, although a smaller amplitude peak appears at ∼1 − 2 ns (Figure 2G). In addition, all constructs show a peak at ∼5 ns of varying amplitudes. This longest-lifetime component overlaps with the donor-only lifetime (Supplementary Figure S3B) and hence corresponds to a component with a FRET close to zero. While there may be some contribution to this so-called “zero-FRET” component from DNA constructs that have a missing or inactive acceptor, our observation that the fractional amplitudes of this component are larger in specific mismatched constructs compared with their matched counterparts and further increase upon Rad4 binding suggests that this component, at least in part, is real and reflects a distorted DNA conformation preferred by Rad4 (5). The lifetime distributions of TTT/TTT show larger amplitudes in the non-B-DNA components, both without and with Rad4, and the fractional amplitudes of these non-B-DNA components increase with increase in temperature (Figure 2H-I). The lifetime distributions for the nonspecific TAT/TAT and the matched ATA/TAT remain relatively unchanged with increase in temperature, and the TAT/TAT distributions are only minimally altered upon Rad4 binding (Figure 2M-O).

**Figure 2.**
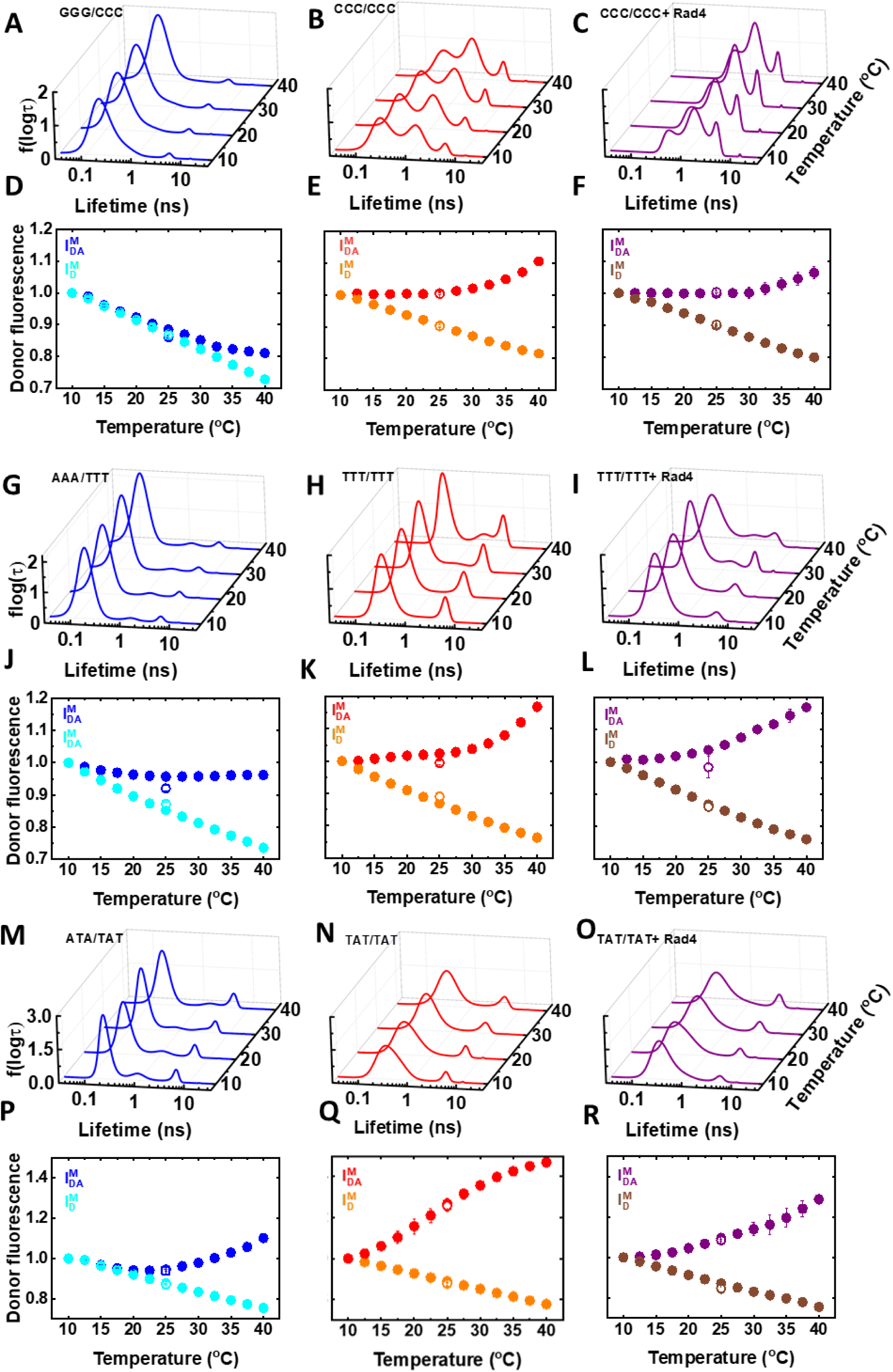
Temperature dependence of fluorescence lifetime distributions and donor fluorescence intensities. Data are shown for the CCC/CCC data set (A-F), the TTT/TTT data set (G-L), and the TAT/TAT data set (M-R). (A-C, G-I, M-O) The fluorescence lifetime distributions are shown for DNA_DA samples at four different temperatures, for matched (blue), mismatched (red), and mismatched+Rad4 (purple). (D-F, J-L, P-R) The corresponding donor fluorescence intensities measured for each of the samples under steady-state conditions in the absence (*I_D_^M^*) and presence (*I_DA_^M^*) of the acceptor are plotted as a function of temperature. The superscript *M* indicates that the measured intensities, *I*_*D*_ and *I*_*DA*_, have been matched at the lowest measured temperature (10 ℃) by dividing by *I*_*D*_(10 ℃) and *I*_*DA*_(10 ℃), respectively. The reversibility of the samples after the heating/cooling cycle was checked by first heating up from 10 to 40 ℃ and then cooling back down to 25 ℃ (open symbols).

The somewhat subtle effects of temperature on matched AAA/TTT, matched ATA/TAT, and mismatched-yet-nonspecific TAT/TAT are readily noted when we plot the temperature dependencies of the donor emission intensities, computed as the area under the measured emission spectra (see SI Methods 1.3 and Supplementary Figure S2), and denoted here as *I*_*D*_ and *I*_*DA*_for DNA_D and DNA_DA samples, respectively. These intensities, when normalized to match at the lowest temperature (10℃), and denoted as *I_D_^M^* and *I_DA_^M^*, respectively, show how the temperature dependencies of *I*_*D*_differ from *I*_*DA*_. In DNA_D samples, the donor intensities *I*_*D*_(or *I_D_^M^*) decrease monotonically with temperature and reflect the decrease in the quantum yield of the donor with increasing temperature (Figures 2D–F, 2J–L, and 2P–R). In contrast, in DNA_DA samples, the donor intensities *I_DA_^M^* exhibit thermal profiles that initially overlap with *I_D_^M^* at low temperatures, but start to deviate at higher temperatures, reflecting changes in FRET E from thermally induced changes in DNA conformations. In the case of the matched GGG/CCC construct, *I_D_^M^* and *I_DA_^M^* are nearly overlapping in almost the entire temperature range measured, with slight deviations noted only above ∼30 ℃ (Figure 2D), equating to essentially no change in FRET with increasing temperature (Figure 2A and Supplementary Figure S1A). In contrast, for AAA/TTT (Figure 2J), ATA/TAT (Figure 2P), and TAT/TAT (Figure 2Q), *I_DA_^M^* starts to deviate from *I_D_^M^* at much lower temperatures, ∼13 ℃, 20 ℃, and 13 ℃, respectively, with the largest deviations over the measured temperature range seen for mismatched TAT/TAT among all the matched or nonspecific constructs (see also Supplementary Figure S1). To summarize the conclusions from the equilibrium measurements, all the matched/nonspecific constructs with As and Ts at the site of interest are more deformable (more readily perturbed with temperature) than the matched GGG/CCC construct. This intrinsic DNA deformability is further enhanced when there are mismatches that are specifically recognized by Rad4, such as CCC/CCC or TTT/TTT (Figures 2E, 2K, and Supplementary Figure S1A–B). Rad4-bound complexes for the specific constructs exhibit distortions in the DNA conformations, as seen in the alterations in the lifetime distributions and the average FRET (Figures 2C, 2I, and Supplementary Figure S1A–B); however, the complex remains dynamic and deformable to nearly the same extent as the specific DNA alone. For the mismatched-yet-nonspecific TAT/TAT, there is no change in the average FRET or the DNA deformability in the presence of Rad4 (Figure 2O and Supplementary Figure S1C), consistent with our results on Rad4 bound to matched DNA constructs reported previously (5).

### Laser T-jump measurements reveal DNA conformational dynamics that span multiple timescales

The DNA conformational distributions of nearly all the constructs we investigated are altered by temperature, making laser T-jump perturbation an appropriate approach to capture their conformational dynamics. In our T-jump apparatus, a 10 ns IR pulse (1550 nm) rapidly heats up a small volume of the sample, the fluorescence of which is probed by a continuous wave light from a Xe lamp, with the excitation wavelength selected at 355 ± 20 nm, to excite the donor (tC°). The tC° probe by itself is insensitive to DNA sequence-context and changes in DNA conformations (12–14); hence, for donor-only (DNA_D) samples, we see only a rapid drop in the donor intensity *I*_*D*_ within the dead-time of our T-jump spectrometer, which is the response of the donor quantum yield from the intensity levels at the initial temperature (*T*_*i*_) to the intensity levels at the final temperature (*T*_*f*_), followed by a slow decay of the donor fluorescence back to the initial (pre-T-jump) intensity levels as the sample temperature cools back, on timescales of 100 − 200 ms (Supplementary Figure S4). Indeed, the size of the T-jump for a given alignment is determined by measurements on donor-only samples (see SI Methods 1.7 and Supplementary Figure S4).

For donor-acceptor (DNA_DA) samples, we start with intensity levels corresponding to *I*_*DA*_(*T*_*i*_) prior to the arrival of the IR pulse. Immediately after the T-jump, we anticipate the same relative drop in donor intensity as in the DNA_D samples, since the temperature response of the donor quantum yield is expected to be independent of the presence of an acceptor in the sample. This drop is illustrated by the black arrow on the equilibrium profile shown in Figure 3A, where the donor fluorescence intensities plotted are normalized with respect to the intensities measured at the initial temperature (and denoted as *I_D_^N^* and *I_DA_^N^* to distinguish them from *I_D_^M^* and *I_DA_^M^*, which denote intensities that are matched at 10 ℃). Next, the donor fluorescence is expected to reach the intensity levels corresponding to the final temperature, *I*_*DA*_(*T*_*f*_), because of DNA conformational changes (illustrated by the purple arrow in Figure 3A and Supplementary Figure S5A), assuming that the entire conformational relaxation is completed prior to ∼50 ms, before the sample temperature itself starts to recover. Finally, the intensity levels are expected to decay back to *I*_*DA*_(*T*_*i*_) on timescales characteristic of when the sample temperature reaches that of the surrounding bath. Therefore, if the entirety of the conformational dynamics were to fall within the T-jump observation window of ∼20 µs to ∼50 ms (i.e., prior to any significant decay of the elevated sample temperature), the amplitude of relaxation kinetics we expect to see (denoted as *Amp*_*eq*_) is proportional to the size of the purple arrow in Figure 3A. The subscript *eq* in *Amp*_*eq*_ is a reminder that these are expected amplitudes, computed from the *equilibrium* profiles. However, not all the predicted amplitudes are, in fact, observed within our T-jump time window, pointing to unresolved or ‘missing’ portions of the relaxation traces, as described below (see also SI Methods 1.10).

**Figure 3.**
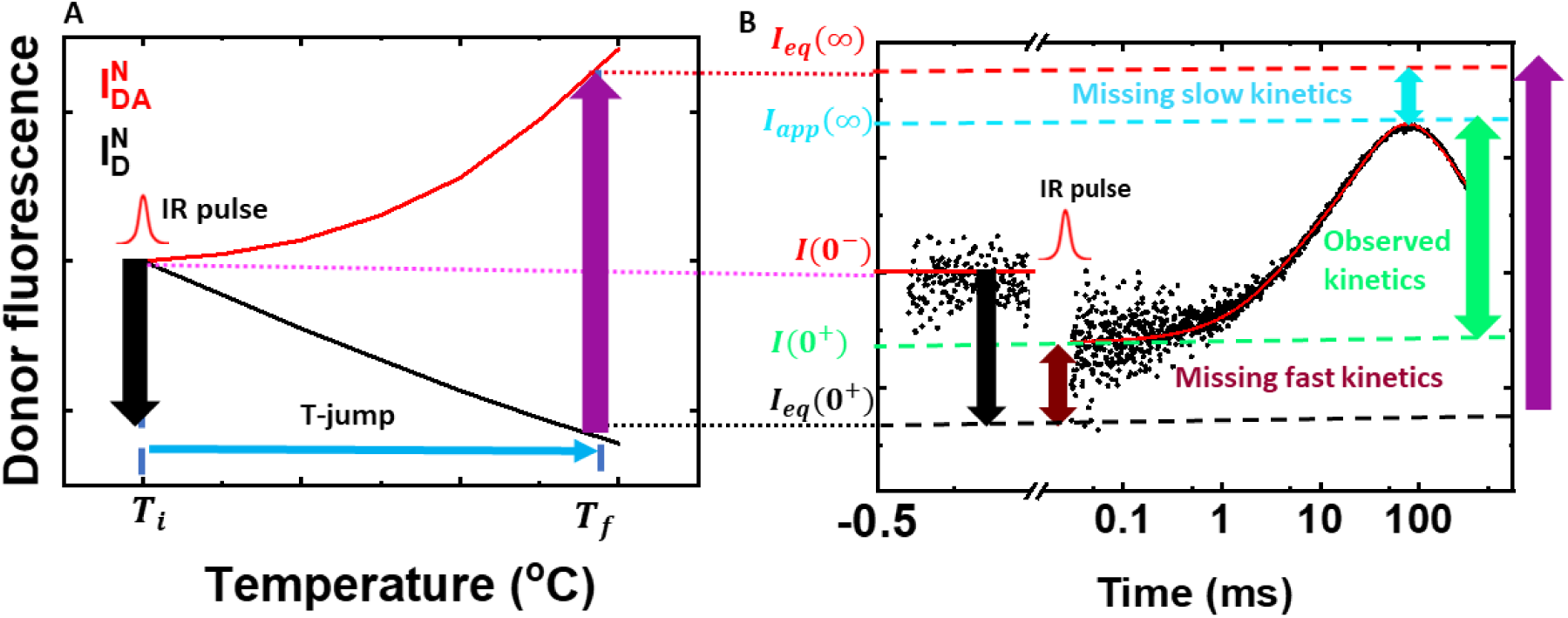
Amplitude analysis from steady-state equilibrium measurements and T-jump traces. (A) Typical thermal profiles of *I*_*D*_ (black) and *I*_*DA*_(red) are shown with the two intensities normalized to match at the initial temperature *T*_*initial*_ (*T*_*i*_). The black vertical arrow pointing down illustrates the decrease in donor intensity for donor-only (DNA_D) samples because of the decrease in the donor quantum yield when the temperature of the sample jumps from *T*_*initial*_ to the final temperature *T*_*final*_ (*T*_*f*_). The purple vertical arrow pointing up illustrates the increase in donor fluorescence expected because of DNA conformational changes in which the DNA conformational distribution shifts from the equilibrium distribution at *T*_*i*_ to that at *T*_*f*_ following the T-jump. This increase in donor fluorescence should appear as conformational relaxation kinetics in a T-jump trace if the timescale of the conformational relaxation falls within the T-jump time window. (B) A representative T-jump trace is shown for a DNA_DA sample. The donor intensity before the arrival of the IR pulse is at the pre-flash level *I*(0^−^), denoted by the red continuous line. The intensity level expected after the T-jump due to donor quantum yield change (computed from the data in panel A) is *I*_*eq*_(0^+^), denoted by the black dashed line. Similarly, the intensity level expected when the conformational population equilibrates at the higher temperature after the T-jump (also computed from the data in panel A) is *I*_*eq*_(∞), denoted by the red dashed line. The measured T-jump trace (black dots) shows that at the earlier recorded time after the IR pulse heats the sample, the donor intensity drops to the level *I*(0^+^) (green dashed line) and then increases to the level *I*_*app*_(∞) (cyan dashed line) before decaying back to the pre-flash level as the temperature in the sample decays back from *T*_*f*_ to *T*_*i*_. The amplitudes of the missing fast, observed, and missing slow kinetics, computed as described in the text, are indicated by the maroon, green, and cyan vertical arrows, respectively.

A typical T-jump trace for a donor-acceptor-labeled (DNA_DA) sample is shown in Figure 3B. The pre-T-jump intensity level is denoted as *I*(0^−^), the intensity level we expect post-T-jump (from purely donor quantum yield changes) is denoted *I*_*eq*_(0^+^), and the intensity level we expect at the end of the conformational relaxation (if the sample temperature could be held indefinitely at the post-T-jump level) is denoted by as *I*_*eq*_(∞) (solid red, dashed black, and dashed red lines, respectively, in Figure 3B and Supplementary Figure S5B). The expected T-jump intensities at the beginning and end of the entire conformational relaxation are readily calculated from the equilibrium intensities as

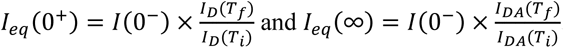

The actual intensity measured at the earliest time point resolved by our T-jump instrument after the arrival of the IR pulse, at ∼20 μs, is *I*(0^+^) (green dashed line in Figure 3B and Supplementary Figure S5B). As seen in the representative T-jump trace, *I*(0^+^) is not as low as *I*_*eq*_(0^+^); we interpret this difference between *I*_*eq*_(0^+^) and *I*(0^+^) as evidence for relaxation kinetics on timescales faster than the ∼20 μs deadtime of our T-jump instrument, and this difference yields the amplitude of the ‘missing’ fast portion of the relaxation trace, denoted as *Amp*_*fast*_(see SI Methods 1.10). On timescales longer than ∼20 μs, we observe relaxation kinetics in which the conformational population distribution shifts from that at the initial temperature (prior to the T-jump) to that at the final temperature (post-T-jump). This relaxation kinetics is characterized by an increase in the donor intensity (because FRET decreases), as anticipated from the equilibrium thermal profiles. As discussed above, if the temperature in the sample could be held at the post-T-jump level, we would expect the donor intensity in DNA_DA samples to eventually reach the computed level *I*_*eq*_(∞); however, the measured intensity levels in the T-jump traces do not always reach that level, if the temperature starts to recover before the conformational relaxation is complete. In that case, the donor intensity reaches a level denoted as *I*_*app*_(∞) (cyan dashed line in Figure 3B and Supplementary Figure S5B) before it starts to decay to the pre-T-jump level *I*(0^−^) with a characteristic time constant of ∼100 − 200 ms for this so-called ‘T-jump recovery’ kinetics.

The *I*(0^+^) and *I*_*app*_(∞) levels for each of the T-jump traces are obtained from MEM analysis of the relaxation traces as described in SI Methods 1.9 and illustrated in Supplementary Figures S6 and S7. The difference between *I*_*eq*_(∞) and *I*_*app*_(∞) is interpreted as evidence for conformational relaxation kinetics that are too slow to be observed within our T-jump time window and yields the amplitude of the ‘missing’ slow portion of the relaxation kinetics denoted as *Amp*_*slow*_. The observed amplitude *Amp*_*obs*_ is then proportional to *I*_*app*_(∞) − *I*(0^+^) (green vertical arrow in Figure 3B) such that *Amp*_*fast*_ + *Amp*_*obs*_ + *Amp*_*slow*_ = *Amp*_*eq*_, the total amplitude expected from the equilibrium measurements. We emphasize here that the clean separation of the two contributions to the full T-jump trace on donor-acceptor samples, from conformational relaxation and from T-jump recovery, is a simplifying approximation because the relaxation is, in fact, coupled to the recovery. However, a more accurate treatment involving temperature-dependent (and therefore time-dependent) relaxation rates, especially on timescales where T-jump recovery dominates, is beyond the scope of this work.

A set of representative relaxation traces for matched, mismatched, and mismatched+Rad4 for each of the constructs are summarized in Supplementary Figure S8–S10. In Supplementary Figure S5, we illustrate the calculations of the expected (*Amp*_*eq*_) and observed (*Amp*_*obs*_) amplitudes for one of these decay traces, and describe how we take into account the variations in these amplitudes from variations in T-jump size when comparing one set of measurements to another. The results of the amplitudes analyses on all the samples measured are summarized in Figure 4.

**Figure 4.**
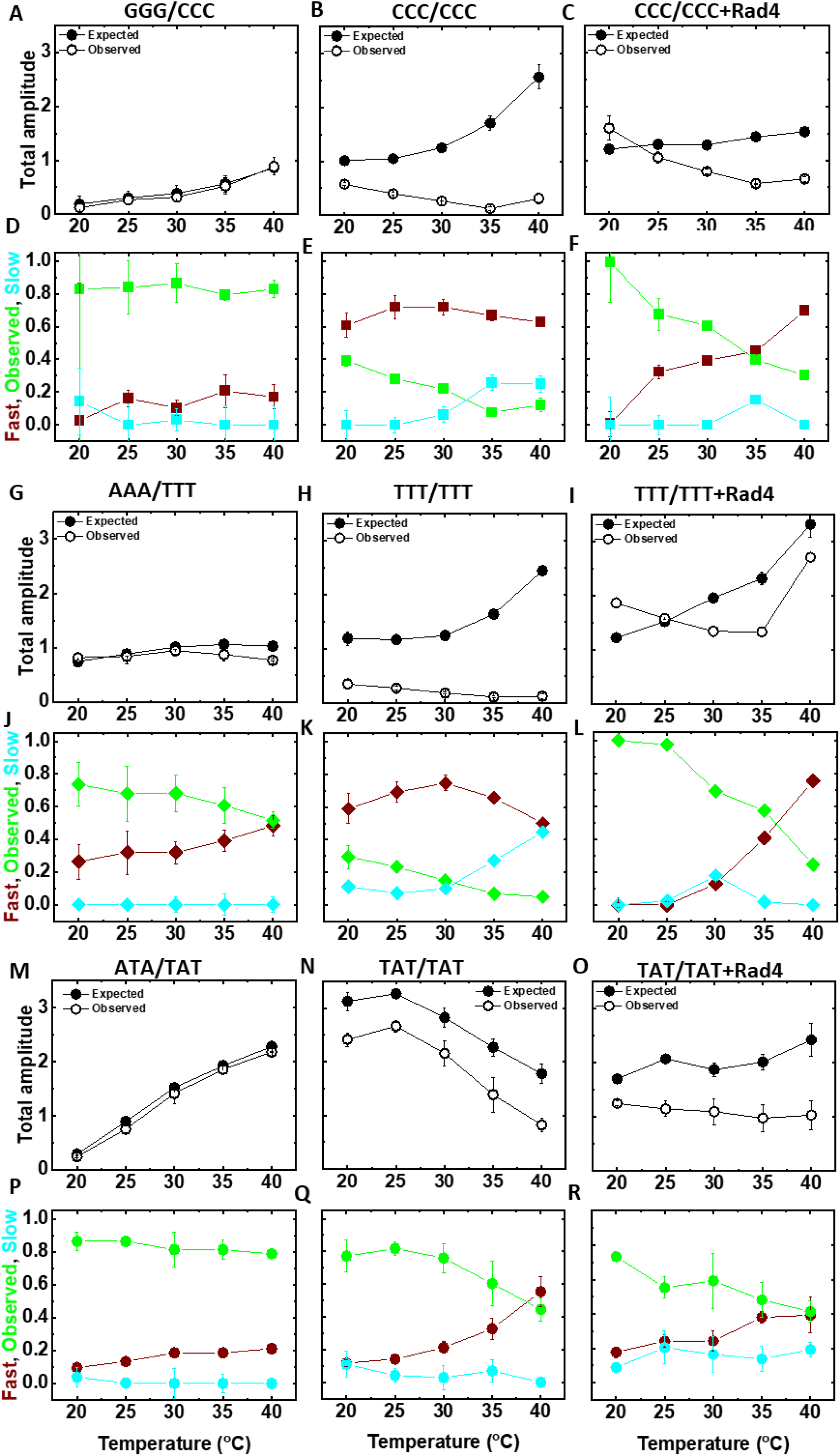
Summary of amplitude analysis results. The expected (from equilibrium; filled circles) and observed (from T-jump; open circles) amplitudes of conformational relaxation— proportional to the purple and green vertical arrows, respectively, in Figure 3B and computed as illustrated in Supplementary Figure S5—are plotted as a function of the final temperature after T-jump for (A-C) the CCC/CCC data set, (G-I) the TTT/TTT data set, and (M-O) the TAT/TAT data set. (D-F, J-L, P-R) The corresponding fractional amplitudes of the missing fast portion (maroon), the observed portion (green), and the missing slow portion (cyan) of the relaxation kinetics are plotted.

### Matched DNA constructs exhibit small amplitude kinetics that fall predominantly in the observable T-jump window

We first examine the relaxation kinetics for the matched constructs. The nearly overlapping equilibrium temperature profiles for the matched donor-only (*I*^*M*^) and donor-acceptor-labeled (*I*^*M*^) for GGG/CCC translate into very little change in FRET with increasing temperature (Figure 2D and Supplementary Figure S1A), indicating a relatively rigid structure. Correspondingly, the conformational relaxation amplitudes from T-jump are also expected to be relatively small (Figure 4A), especially at the low temperature end, but increase as the sample temperature is raised. The measured amplitudes on GGG/CCC constructs are in excellent agreement with these predictions, i.e., we detect small amplitude relaxation dynamics that fall within our T-jump time window, with little or no ‘missing’ kinetics (Figure 4D and Supplementary Figure S8D, G).

In contrast, matched AAA/TTT and ATA/TAT exhibit equilibrium temperature profiles in which *I_D_^M^* and *I_DA_^M^* diverge as the temperature is raised, indicating a comparatively more deformable DNA structure that is readily perturbed with temperature increase (Figure 2J, 2P, and Supplementary Figure S1B, C). The expected T-jump relaxation amplitudes are correspondingly larger than for GGG/CCC (Figure 4G, M). The observed relaxation amplitudes fall slightly short of the expected amplitudes, especially at final temperatures above ∼30 ℃, and the difference appears as a missing fast phase that ranges from ∼25% of the total expected amplitude at 20 ℃ to ∼50% at 40 ℃ for AAA/TTT and from ∼10% at 20 ℃ to ∼20% at 40 ℃ for ATA/TAT (Figure 4J, P). Neither of these matched constructs exhibit evidence for any missing slow phase in the temperature range measured. We conclude that, compared to GGG/CCC, the AT-rich AAA/TTT and ATA/TAT exhibit relatively larger amplitude motions with some fraction of the dynamics occurring on sub-20 μs timescale. These results showcase the remarkable sensitivity of our T-jump apparatus to detect small amplitude DNA conformational dynamics (unwinding and bending motions) and are the first such measurements on matched DNA.

### Mismatched-but-nonspecific TAT/TAT exhibits kinetics that mimics AT-rich matched DNA

Next, we look at the relaxation kinetics of mismatched TAT/TAT, which is not recognized by Rad4 as a specific substrate. Here too, the relaxation kinetics lie predominantly within the observable T-jump window, especially at temperatures below ∼30 ℃ (Figure 4N, Q and Supplementary Figure S10E, H). Above that temperature, the missing fast phase appears, with its amplitude increasing up to ∼55% at 40 ℃. This construct also shows some evidence for missing slow kinetics, although with less than 10% amplitude (Figure 4Q). Taken together, we conclude that all AT-rich constructs, whether matched or mismatched-but-nonspecific, exhibit similar conformational relaxation kinetics, with amplitudes significantly larger than those seen in the matched GGG/CCC.

### Mismatched-specific (CCC and TTT) exhibit significantly larger amplitudes in missing fast and slow kinetics

In Supplementary Figure S11, we compare the expected amplitudes from a T-jump perturbation for matched versus mismatched constructs, which reflects the change in FRET when the temperature is raised and is a gauge of the deformability of that DNA construct. We conclude that, while all mismatched DNA are more deformable/dynamic than their matched counterparts, the dynamism by itself is not a good predictor of which mismatched site is specifically recognized by Rad4. For example, the nonspecific TAT/TAT is predicted to have the largest amplitude for relaxation dynamics, especially at the low temperature end, when compared with the specific CCC/CCC and TTT/TTT constructs. However, what separates the specific from the nonspecific constructs is the extent to which the observed relaxation amplitudes (the fraction that falls within the T-jump window) differs from the expected amplitudes. For both the specific constructs, nearly 60-80% of the total expected amplitude is missing at the fast end of the relaxation kinetics over the temperature range measured (Figures 4B, E and 4H, K). Additionally, ∼30–40% of the total expected amplitude is missing at the slow end of the relaxation kinetics for both these constructs at temperatures ≥ 35 ℃. In contrast, the mismatched-yet-nonspecific TAT/TAT exhibited very low-level missing amplitudes at the slow end, and the matched constructs showed none. We conclude that what distinguishes the specific from the nonspecific constructs is large amplitude rapid motions (on timescales < 20 μs) as well as much slower dynamics (on timescales > 50 ms); the slow dynamics grow in amplitude as the temperature is raised. With Rad4 bound to specific DNA, the missing fast amplitude is suppressed at lower temperatures and the missing slow amplitude is significantly suppressed (Figures 4F, L).

### Observed conformational dynamics span more than a decade in time

For all constructs that showed significant dynamics with T-jump perturbation, the relaxation kinetics within the observed T-jump window could not be described as a single-exponential decay, as evident from the broad distribution of relaxation times obtained from MEM analysis of the T-jump traces (Figure 5). These analyses indicate at least two distinct peaks in the MEM distributions, suggesting a minimum of two exponentials needed to describe the observed relaxation traces, in addition to the fast and slow portions that lie outside the observed window. These conformational dynamics likely represent a combination of spontaneous DNA motions that span several timescales such as base-pair breathing (local melting), DNA twisting/unwinding, DNA bending, and base flipping, all of which are potentially detected by the tC° and tC_nitro_ FRET probes and showcase the sensitivity of this FRET pair.

**Figure 5.**
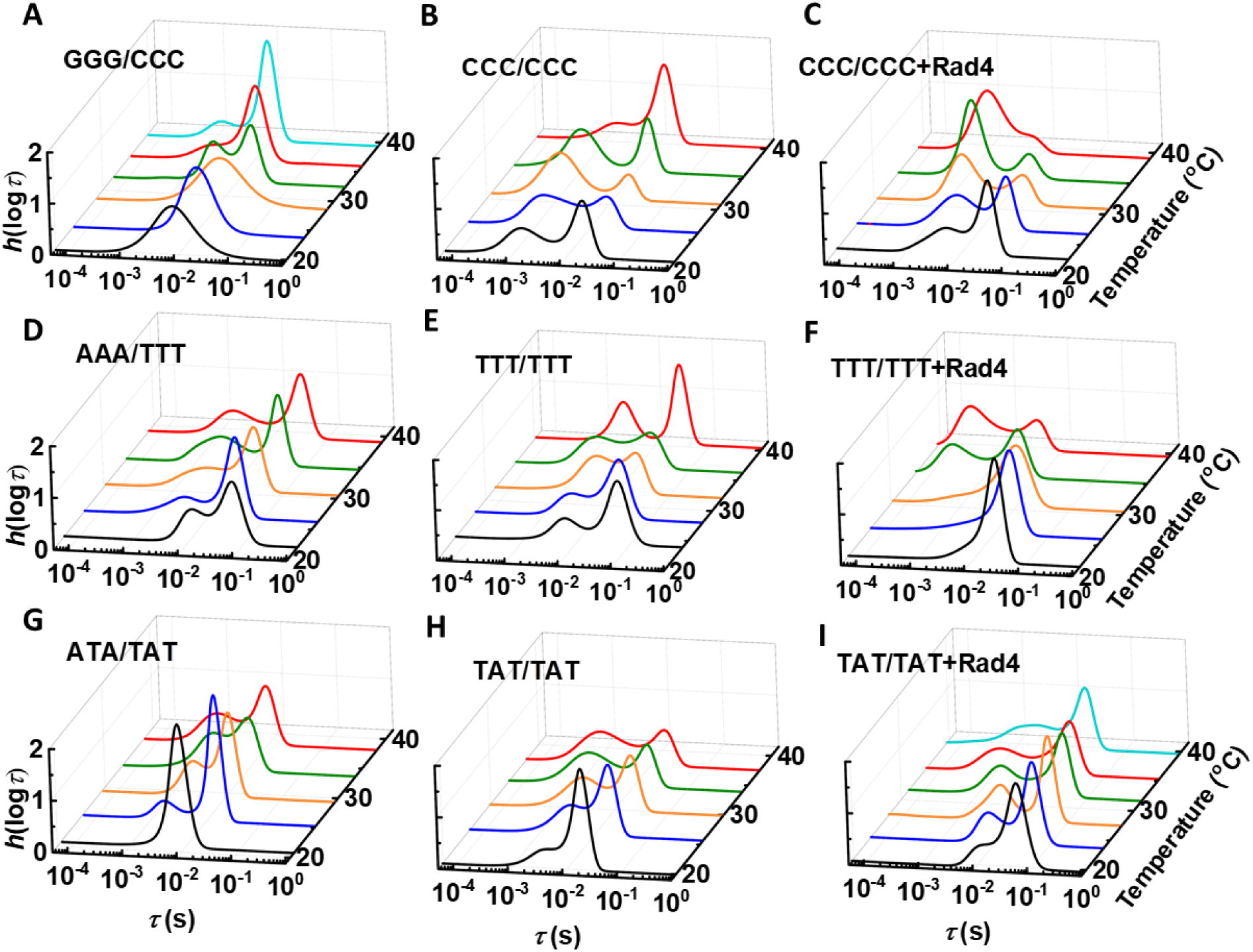
Distribution of relaxation rates from MEM analysis of T-jump traces. MEM distributions of the relaxation times ℎ(logτ) that describe the observed conformational relaxation kinetics are plotted versus the relaxation time τ for (A-C) the CCC/CCC data set; (D-F) the TTT data set; and (G-I) the TAT/TAT data set. All distributions are renormalized such that the area under the curve is 1.

## DISCUSSION

### Conformational dynamics in matched and mismatched DNA are probed by time-resolved FRET measurements with tC°-tC_nitro_ probes

In this study, we sought to investigate the nature of intrinsic fluctuations in a series of DNA oligomers with 3 base-pair mismatches that were previously shown to be excellent model systems for understanding the mechanism of damage sensing by the NER protein Rad4/XPC. Depending on the nature of the mismatch, these constructs are recognized by Rad4 with high (CCC/CCC), low (TTT/TTT), or no (TAT/TAT) specificity. In a previous study, we showed that the CCC/CCC mismatch exhibited a range of DNA conformations that, on average, deviated significantly from a canonical B-DNA conformation; furthermore, the amplitude of the distorted conformations increased with temperature (5). In direct contrast, the matched GGG/CCC construct exhibited predominantly B-DNA conformation that remained largely unaltered with increasing temperature. The intermediate construct TTT/TTT and nonspecific TAT/TAT showed deviations from B-DNA but these deviations were significantly smaller than those observed in CCC/CCC. These earlier studies were performed with fluorescence lifetime measurements that provided a snapshot of the accessible conformations at each temperature but gave no insights into the timescales of these fluctuations. However, the implication from these snapshots was that mismatched constructs recognized efficiently by Rad4 are the ones that are intrinsically more dynamic and more readily deformed. The two features are expected to go hand-in-hand, since more dynamic DNA sequences are the ones that likely have weaker stacking interactions, which also renders them more amenable to being unwound and distorted. This intrinsic deformability implies that Rad4 can more readily flip out nucleotides from the DNA duplex to form the so-called ‘open’ conformation seen in all the crystal structures of Rad4 bound to specific DNA sites (or tethered to nonspecific sites). More dynamic DNA sequences are also implicated in modifying the interactions of Rad4/XPC with DNA, thereby altering the free energy landscape for a diffusing protein in search of damage sites, and perhaps triggering a conformational change in the protein that prevents it from diffusing away from the seemingly anomalous site (8).

To see what features distinguish the specific from the nonspecific DNA substrates, we herein examined the amplitudes and timescales of conformational changes induced by a laser T-jump perturbation. Conformational dynamics were observed in nearly all the constructs; in general, mismatched DNA exhibited larger amplitude dynamics than the matched counterparts. The exception was the matched GGG/CCC construct that showed negligible amplitude dynamics at 20 ℃, and which became significant only at 35 − 40 ℃ range, indicating a relatively rigid structure that resisted perturbation with increase in temperature. All other constructs, specific or nonspecific, were readily perturbed when the temperature was raised, and exhibited conformational relaxation dynamics over multiple timescales, spanning not just the time window of our T-jump spectrometer (∼20 µs – 50 ms) but also detected as “missing” portions outside the T-jump window, including fast kinetics (dynamics on < 20 µs timescale) and in some cases slow kinetics (dynamics on > 50 ms timescale). Below we expand on these results and reiterate the subtle though important variations between the specific and the nonspecific constructs.

### Nonspecific constructs with As and Ts at the site of interest were more dynamic than those with Gs and Cs

The matched GGG/CCC was the most rigid (least dynamic) of all the nonspecific constructs that we looked at. This was also the only nonspecific construct we studied that had Gs and Cs at the site of interest, the other three being ATA/TAT, TAT/TAT, and AAA/TTT. The larger amplitude fluctuations detected in A/T rich sites, whether matched or mismatched, are consistent with the higher propensity of these pairs to transiently melt (15–17). The actual nature of DNA motions observed in our measurements is not evident from our study, but given the sensitivity of our probes to unwinding fluctuations, it is not unreasonable to conclude that breathing dynamics that transiently melt a short stretch of DNA would amplify the twisting/unwinding fluctuations. The G/C rich sites of matched sequences are less prone to such melting, and accordingly, the conformational dynamics detected by our probes in the context of GGG/CCC were suppressed.

One intriguing observation from this study is that the trends exhibited by the nonspecific AT-rich constructs did not follow any simple pattern with increase in temperature. For AAA/TTT, the total amplitudes expected from T-jump perturbation remained relatively unchanged as the temperature of the sample was raised (Figure 4G); for ATA/TAT, these amplitudes increased with increase in temperature (Figure 4M); for the mismatched-yet-nonspecific TAT/TAT, they decreased with increase in temperature (Figure 4N). Note that the highest temperature in our kinetics measurements (∼40 ℃ final temperature, after T-jump) was well below the overall melting temperatures of these duplexes, which were measured by absorbance melting profiles to be in the range of ∼60 − 80 ℃ (see table in Supplementary Figure S1A). The FRET measurements, however, report on local unwinding/melting. A plausible explanation for these observed trends in the T-jump detected amplitudes is as follows. The ATA/TAT site is relatively intact at 10 ℃ and requires temperatures above ∼20 ℃ to begin to be perturbed, as evidenced by overlapping profiles of the normalized donor intensities *I_D_^N^* and *I_DA_^N^* up to at least 20 ℃ (Figure 2P), and small amplitude dynamics at low temperatures that increase dramatically as the temperature increases, likely from promotion of unstacking/unpairing at the elevated temperatures (Figure 4M). In contrast, TAT/TAT appears already partially unpaired at 10 ℃, as evidenced by how readily it is perturbed as the temperature is raised (*I_D_^N^* and *I_DA_^N^* show diverging behavior starting at the lowest temperature; Figure 2Q), with further increase in temperature having a diminishing effect. This behavior is reflected in substantial amplitude in T-jump expected/observed dynamics at the low-temperature end, which steadily decreases as the temperature is further raised and the site is completely disrupted (Figure 4N). The AAA/TTT site is in some sense an outlier in that the total expected/observed amplitude remained relatively unchanged with increase in temperature (Figure 4G). This lack of temperature dependence in the observed amplitude suggests that the origin of the conformational changes induced by T-jump for this site are not related to local untacking/melting but rather reflect a conformational switch between two intact but competing structures. This behavior of the AAA/TTT construct may be a consequence of the unusual properties of A-tracts, typically defined as three or more A bases in a row, which have been reported to be relatively rigid in some respects, but also unusually flexible in other aspects, and whose deformability and dynamics remain puzzling (18,19).

### Conformational dynamics in mismatched constructs specifically recognized by Rad4 are distinct from those in nonspecific constructs

The nonspecific AT-rich constructs (ATA/TAT, TAT/TAT, AAA/TTT) exhibited conformational relaxation kinetics predominantly within the T-jump window, with typically less than 20% of the total expected amplitude at 20 ℃ missing (or unresolved) at the fast end (i.e. faster than the ∼20 µs dead time of our T-jump spectrometer). This amplitude in the fast/unresolved kinetics increased to at most ∼50% at 40 ℃ for two of the three (AAA/TTT and TAT/TAT). Notably, none of the nonspecific constructs showed significant evidence for missing amplitude at the slow end of the observed kinetics. We note here that the mismatched-yet-nonspecific TAT/TAT did exhibit low (< 10%) amplitude in the missing slow kinetics, but there was no real trend with temperature. The specific constructs CCC/CCC and TTT/TTT were distinct from the nonspecific constructs in two aspects. First, the amplitudes in the missing fast portion were significantly larger in both CCC/CCC and TTT/TTT, more than 50% within the entire temperature range and as high as 70–80% near 25 − 30 ℃. Second, the specific constructs alone exhibited clear trends for amplitudes missing at the slow end, which appeared above ∼30 ℃ and increased in their relative fractions as the temperature was raised, up to 30–40% at 40 ℃.

### Conformational dynamics measured with tC°-tC_nitro_ FRET pair span multiple timescales

In an earlier study, we examined equilibrium fluctuations in CCC/CCC and GGG/CCC constructs using confocal microscopy-based fluorescence correlation spectroscopy (FCS) on freely diffusing DNA oligomers (20). These measurements, done using Atto-dyes as a FRET pair attached to the DNA with linkers, detected fluctuations in the CCC/CCC construct but not in GGG/CCC, consistent with the current results. The FCS measurements revealed the timescales of these dynamics to overlap with diffusion times of the oligomers through the confocal volume; deconvolution of the diffusion contribution to the FCS correlation function resulted in the correlation function attributable to conformational dynamics that could be well described as a single exponential decay with a characteristic relaxation time of ∼300 μs at 20 ℃. The conformational dynamics detected in this study span many orders of magnitude in time and reflect the sensitivity of both the T-jump apparatus and the tC°-tC_nitro_ FRET pair to capture even small amplitude motions in DNA with unprecedented time resolution and sensitivity.

## CONCLUSIONS

In this study we leveraged the sensitivity of laser T-jump spectroscopy with the unique properties of the cytosine analog FRET pair tC° and tC_nitro_ to detect subtle changes in DNA conformations in response to a T-jump perturbation. Using a novel and rigorous ‘amplitude analysis’ approach, we unveiled conformational dynamics in both matched and mismatched DNA over a broad range of timescales, including evidence of missing fast or slow kinetics that fell outside the T-jump-resolved observation window of ∼20 μs − 50 ms. Together with insights gained from previous lifetime studies, we find that GC-rich matched sites exhibit low amplitude fluctuations, consistent with more rigidly stacked and strongly paired sites, while AT-rich sites exhibit larger amplitude fluctuations over many timescales that range from sub-20 μs up to ∼50 ms. The dynamic range of these motions increased when there were mismatches present. Most important, 3-bp mismatched sites specifically recognized by Rad4 (CCC/CCC and TTT/TTT) also showed evidence of slow (> 50 ms) dynamics, the amplitudes of which increased with increasing temperature. We conclude that these slow dynamics reflect fluctuations to more distorted DNA conformations that present a higher free energy barrier, but nonetheless a barrier that these DNA constructs are capable of overcoming given enough thermal energy. Mismatched-but-nonspecific DNA did not exhibit these slow dynamics up to 40 ℃. We conclude that Rad4 shows an affinity for those DNA sites that can spontaneously access the distorted conformations that Rad4 prefers, albeit with a higher free energy barrier than if Rad4 was assisting in achieving those distortions. These conclusions are not inconsistent with a conformational capture mechanism in damage sensing, but rather suggest that there are multiple pathways by which the protein and DNA fall into the free energy well of the recognition complex. This conclusion is also in line with the kinetic gating mechanism in that if the free energy barrier for forming the ‘open’ recognition complex is too high, Rad4 will bypass that site in search of a more pliable one, more amenable to being unwound.

## ACKNOWLEDGEMENTS

This work was supported by grants from the National Science Foundation (MCB-1715649 and MCB-2107527 to A.A. and MCB 2131806 to J.-H. M) and the National Institutes of Health (GM147899 to J.-H. M). P.J.S acknowledges support from the Office of Science Management and Operations (OSMO) of the National Institute of Allergy and Infectious Diseases, National Institutes of Health.

## Notes

### Competing Interest Statement

The authors have declared no competing interest.

### Summary of Updates

The manuscript has been revised to adapt the abstract for the journal to which the manuscript will be submitted and to update some details of the analysis presented.

## REFERENCES

1. Min, J.H. and Pavletich, N.P. (2007) Recognition of DNA damage by the Rad4 nucleotide excision repair protein. Nature, 449, 570–575.

2. Paul, D., Mu, H., Zhao, H., Ouerfelli, O., Jeffrey, P.D., Broyde, S. and Min, J.H. (2019) Structure and mechanism of pyrimidine-pyrimidone (6-4) photoproduct recognition by the Rad4/XPC nucleotide excision repair complex. Nucleic Acids Res, 47, 6015–6028.

3. Chen, X., Velmurugu, Y., Zheng, G., Park, B., Shim, Y., Kim, Y., Liu, L., Van Houten, B., He, C., Ansari, A., et al. (2015) Kinetic gating mechanism of DNA damage recognition by Rad4/XPC. Nat Commun, 6, 5849.

4. Velmurugu, Y., Chen, X., Slogoff Sevilla, P., Min, J.H. and Ansari, A. (2016) Twist-open mechanism of DNA damage recognition by the Rad4/XPC nucleotide excision repair complex. Proc Natl Acad Sci U S A, 113, E2296–2305.

5. Chakraborty, S., Steinbach, P.J., Paul, D., Mu, H., Broyde, S., Min, J.H. and Ansari, A. (2018) Enhanced spontaneous DNA twisting/bending fluctuations unveiled by fluorescence lifetime distributions promote mismatch recognition by the Rad4 nucleotide excision repair complex. Nucleic Acids Res, 46, 1240–1255.

6. Paul, D., Mu, H., Tavakoli, A., Dai, Q., Chakraborty, S., He, C., Ansari, A., Broyde, S. and Min, J.H. (2021) Impact of DNA sequences on DNA ’opening’ by the Rad4/XPC nucleotide excision repair complex. DNA Repair (Amst), 107, 103194.

7. Kong, M., Liu, L., Chen, X., Driscoll, K.I., Mao, P., Bohm, S., Kad, N.M., Watkins, S.C., Bernstein, K.A., Wyrick, J.J. et al. (2016) Single-molecule imaging reveals that Rad4 employs a dynamic DNA damage recognition process. Mol Cell, 64, 376–387.

8. Cheon, N.Y., Kim, H.S., Yeo, J.E., Scharer, O.D. and Lee, J.Y. (2019) Single-molecule visualization reveals the damage search mechanism for the human NER protein XPC-RAD23B. Nucleic Acids Res, 47, 8337–8347.

9. Steinbach, P.J., Ionescu, R. and Matthews, C.R. (2002) Analysis of kinetics using a hybrid maximum-entropy/nonlinear-least-squares method: application to protein folding. Biophys. J., 82, 2244–2255.

10. Sternisha, S.M., Whittington, A.C., Martinez Fiesco, J.A., Porter, C., McCray, M.M., Logan, T., Olivieri, C., Veglia, G., Steinbach, P.J. and Miller, B.G. (2020) Nanosecond-Timescale Dynamics and Conformational Heterogeneity in Human GCK Regulation and Disease. Biophys J, 118, 1109–1118.

11. Preus, S., Kilsa, K., Miannay, F.A., Albinsson, B. and Wilhelmsson, L.M. (2013) FRETmatrix: a general methodology for the simulation and analysis of FRET in nucleic acids. Nucleic Acids Res, 41, e18.

12. Sandin, P., Wilhelmsson, L.M., Lincoln, P., Powers, V.E., Brown, T. and Albinsson, B. (2005) Fluorescent properties of DNA base analogue tC upon incorporation into DNA--negligible influence of neighbouring bases on fluorescence quantum yield. Nucleic Acids Res, 33, 5019–5025.

13. Sandin, P., Borjesson, K., Li, H., Martensson, J., Brown, T., Wilhelmsson, L.M. and Albinsson, B. (2008) Characterization and use of an unprecedentedly bright and structurally non-perturbing fluorescent DNA base analogue. Nucleic Acids Res, 36, 157–167.

14. Wilhelmsson, L.M., Holmen, A., Lincoln, P., Nielsen, P.E. and Norden, B. (2001) A highly fluorescent DNA base analogue that forms Watson-Crick base pairs with guanine. J Am Chem Soc, 123, 2434–2435.

15. Altan-Bonnet, G., Libchaber, A. and Krichevsky, O. (2003) Bubble dynamics in double-stranded DNA. Phys. Rev. Lett., 90, 138101.

16. Ambjornsson, T., Banik, S.K., Krichevsky, O. and Metzler, R. (2007) Breathing dynamics in heteropolymer DNA. Biophys J, 92, 2674–2684.

17. Zeida, A., Machado, M.R., Dans, P.D. and Pantano, S. (2012) Breathing, bubbling, and bending: DNA flexibility from multimicrosecond simulations. Phys Rev E Stat Nonlin Soft Matter Phys, 86, 021903.

18. Johnson, S., Chen, Y.J. and Phillips, R. (2013) Poly(dA:dT)-rich DNAs are highly flexible in the context of DNA looping. PLoS One, 8, e75799.

19. Marin-Gonzalez, A., Vilhena, J.G., Perez, R. and Moreno-Herrero, F. (2021) A molecular view of DNA flexibility. Q Rev Biophys, 54, e8.

20. Ten, T.B., Zvoda, V., Sarangi, M.K., Kuznetsov, S.V. and Ansari, A. (2022) “Flexible hinge” dynamics in mismatched DNA revealed by fluorescence correlation spectroscopy. J. Biol. Phys., 48, 253–272.

