## Supporting Information for "Evidence for intrinsic DNA dynamics and deformability in damage sensing by the Rad4/XPC nucleotide excision repair complex"

### SI Methods

**1.1 Preparation of double-stranded DNA substrates.** The concentrations of each single-stranded oligonucleotide (purchased from TriLink Biotechnologies) were determined by UV absorbances at 260 nm using the extinction coefficients provided by TriLink Biotechnologies, based on the nearest neighbor method (1,2). The errors in the measured DNA concentrations were estimated to be less than ~10%. To prepare duplex DNA, two complementary oligonucleotides, each at 100 µM, were mixed in phosphate-buffered saline (PBS) buffer (10 mM disodium hydrogen phosphate ($Na_2HPO_4$), 2 mM monopotassium phosphate ($KH_2PO_4$), 137 mM NaCl, 2.7 mM potassium chloride (KCl), pH 7.4) in a microcentrifuge tube. The tube was then immersed in 2 L of ~100 °C water bath on a hot plate. After 10 min, the hot plate was turned off and the oligonucleotides were annealed as the water bath was cooled down slowly to room temperature over 5 to 6 hours.

**1.2 Preparation of Rad4–Rad23 complexes.** All Rad4-Rad23 complexes were prepared as previously described (3,4). Briefly, Hi5 insect cells co-expressing Rad4 and Rad23 proteins were harvested 2 days after infection. After lysis, the protein complex was purified by affinity chromatography (Ni-NTA Agarose, MCLAB), anion-exchange (Source Q, GE healthcare) and cation exchange (Source S, GE healthcare) chromatography followed by gel-filtration (Superdex 200, GE healthcare), and further concentrated by ultrafiltration to ~13–14 mg/ml (135–150 µM) in 5 mM bis-tris propane–HCl (BTP-HCl), 800 mM NaCl and 5 mM DTT, pH 6.8, for all constructs used in this study.

**1.3 Equilibrium FRET from steady-state measurements.** The fluorescence measurements under equilibrium conditions were carried out on 10 µM $tC^o/tC_{nitro}$-labeled DNA in the presence and absence of equimolar (10 µM) proteins in PBS with 1 mM DTT. The fluorescence data were acquired by a FluoroMax4 spectrofluorometer (JobinYvon, Inc., NJ). The raw data for each sample consists of fluorescence emission spectra measured in the wavelength range 375–550 nm, with excitation of the donor ($tC^o$) at 365 nm; the spectra were measured over the temperature range of 10 °C to 40 °C in 2.5 °C increments. Corresponding measurements of buffer-only control samples were done under identical conditions. To ensure the integrity of each sample after heating, the samples were cooled back down to 25 °C and the fluorescence intensities were measured again at that temperature.

The FRET efficiency ($E$) was obtained using the 'donor fluorescence ratio' method (5), in which the decrease in the donor tC° fluorescence in the presence of acceptor $tC_{nitro}$ compared with its absence is a measure of the FRET efficiency between the pair (6). The donor fluorescence emission intensities were measured in two separate samples: $I_D$ from the donor-only sample (DNA_D) and $I_{DA}$ from the donor-acceptor sample (DNA_DA), and the FRET efficiency was calculated as $E = 1 - \frac{I_{DA}}{I_D}$. For each sample, the characteristic emission intensities $I_D$ and $I_{DA}$ were computed as the area under the corresponding emission curves in the range of 425–475 nm. The error bars are the standard error of the mean (s.e.m.) from 2-6 independent sets of measurements.

For the temperature scan results depicting fluorescence intensities $I_D^M$ (or $I_{DA}^M$) versus temperature, each scan was normalized to match the $I_D$ (or $I_{DA}$) intensities at the lowest temperature (at 10 °C) prior to computing the mean and s.e.m values from the independent sets of measurements. Similary, for the purpose of amplitude analysis (see SI Methods 1.10), the measured fluorescence intensities were matched at the initial temperature ($T_i$) to obtain $I_D^N$ (or $I_{DA}^N$) and the corresponding intensities at the final temperature ($T_f$) and the associated errors were recomputed, as described in SI Methods 1.11.

The 'donor ratio' method requires ratio of fluorescence intensities from two separate samples and any errors in matching the concentrations of the two samples appear as a systematic error in the calculation of the FRET efficiencies. Fluorescence lifetime measurements (described below) are unaffected by variations in the sample concentrations and hence provide a more accurate determination of FRET efficiencies. The FRET values from steady-state measurements were therefore adjusted to match those from fluorescence lifetime measurements at 20 °C, prior to the comparison of the temperature-dependence of FRET from both methods, shown in Supplementary Figure S1.

**1.4 Equilibrium FRET from lifetime measurements.** Fluorescence decay curves were measured with a PicoMaster fluorescence lifetime instrument (HORIBA-PTI, London, Ontario, Canada). The excitation source was a Fianium Whitelase Supercontinuum laser system (maximum power 4W), which produces ~6 ps broad band pulses. For excitation of tC°, the laser pulses were passed through a monochromator set at 365 nm (bandpass 10 nm) followed by a 355/40 nm Semrock BrightLine® single-band bandpass filter. The emission from the sample was collected orthogonal to the excitation beam after passing through a 470/100 nm Semrock BrightLine® single-band bandpass filter followed by another monochromator set at 460 nm (bandpass 10 nm) and detected by a Hamamatsu microchannel plate photomultiplier (MCP-650). The instrument response function (IRF) of the system was measured using a dilute aqueous solution of Ludox (Sigma-Aldrich). The full width at half maximum (fwhm) of the IRF was ~100 ps. Fluorescence decay curves were recorded on a 100 ns timescale, resolved into 4096 channels, to a total of 10,000 counts in the peak channel, with the repetition rate of the laser adjusted to 10 MHz.

For all FRET measurements, decay traces were measured for donor-only ($tC^o$-labeled) duplexes without acceptor ($tC_{nitro}$) as well as donor-acceptor ($tC^o$-$tC_{nitro}$-labeled) duplexes. The decay traces were analyzed using the maximum entropy method (MEM), as described previously (7), to obtain an effective discretized distribution of log-lifetimes $f_j(log\tau_j)$ for the $j$-th lifetime $\tau_j = 10^{\log\tau_j}$ (normalized such that $\sum_j f_j\, d(\log\tau_j) = 1$). For donor-only samples, the MEM outputs gave narrow distributions consistent with single-exponential decays. For donor-acceptor-labeled samples, the distributions were broader. The average FRET efficiency for each sample was computed as $<E>= 1 - \frac{<\tau_{DA}>}{<\tau_D>}$, where $<\tau_D>= \sum_j f_j \tau_{Dj} d(\log\tau_j)$ and $<\tau_{DA}>= \sum_j f_j \tau_{DAj} d(\log\tau_j)$ are the average donor lifetimes (averaged over the entire distribution) in the absence and presence, respectively, of the acceptor.

**1.5 Laser T-jump spectrometer.** Kinetic measurements were carried out using a home-built laser temperature jump (T-jump) apparatus, which uses 10-ns laser pulses at 1550 nm, generated by Raman shifting the 1064 nm pulses from the output of an Nd:YAG laser, to rapidly heat a small volume of the sample within ~10 ns (8). The laser pulses were focused to ~1 mm spot size onto a 2-mm wide sample cuvette of path length 0.5 mm. Each laser pulse (~40 mJ/pulse at the sample position) yielded ~4–10 °C T-jump at the center of the heated volume. The laser was operated at a repetition rate of 1 Hz, to give enough time between pulses for the T-jump to recover back to the initial temperature before the arrival of the next pulse.

The probe source for excitation of $tC^o$-labeled samples was a 200-W Hg-Xe lamp, with the excitation wavelengths selected by a broadband filter (355/40 BrightLine filter, Semrock, Rochester, New York). The fluorescence emission intensity was monitored using a Hamamatsu

R928 photomultiplier tube equipped with another broadband filter (470/100 BrightLine filter, Semrock, Rochester, New York) and a 500 MHz transient digitizer (Tektronix, DPO4054B).

The initial temperature of the sample was measured using a thermistor (Precision Epoxy NTC, 44008RC) in direct contact with the sample cell, the temperature of which was controlled by a heat bath (Neslab, TRE111). The magnitude of the temperature change upon T-jump perturbation was determined from measurements on control (donor-only) samples in which no conformational relaxation kinetics are expected, as described in SI Methods 1.7).

**1.6 Acquisition and analyses of T-jump relaxation traces.** We typically acquired the kinetics traces on three timescales, with one million data points in each trace. The shortest timescale covered kinetics up to 1.6 ms with a time-resolution of 1.6 ns; the intermediate timescale covered up to 32 ms with a time-resolution of 32 ns; the longest timescale covered up to 320 ms with a time-resolution of 320 ns. For each timescale, 512 kinetics traces were acquired and averaged by the digitizer and saved for further analysis. The temporal data acquired on a linear time grid was subsequently averaged to 100 points per logarithmic decade. This logarithmic averaging preserved high temporal resolution at the shorter timescales (at the expense of larger errors) and allowed more data points to be averaged at longer times (thereby reducing the errors at the longer times). Kinetic traces acquired over the three different timescales were matched and combined into a single trace for further analysis, as described in SI Methods 1.8. Prior to combining the different traces, data acquired below ~20 μs in each trace were discarded because of artifacts either from scattered infrared laser light into the photomultiplier tube, or due to cavitation effects from microbubbles in the samples (9). Thus, the deadtime of our instrument is ~20 μs.

**1.7 Measurements of T-jump size and characteristic time for T-jump decay.** The magnitude of T-jump for a given alignment of the T-jump spectrometer was determined by measurements on donor-only control samples (Supplementary Figure S4). Steady-state measurements on donor-only samples showed a monotonic decrease in the donor fluorescence intensity that reflected a decrease in the donor quantum yield with increase in temperature (Supplementary Figure S4A). Correspondingly, T-jump measurements on donor-only samples showed a drop in the donor intensity from the pre-T-jump level (denoted as $I(0^{-})$) to the post-T-jump level (denoted as $I(0^{+})$), which eventually decayed to the pre-T-jump level as the temperature of the sample decayed back to that of the surrounding bath (Supplementary Figure

S4B). The corresponding intensities in the steady-state measurements would be $I_D(T_i)$ and $I_D(T_f)$, as illustrated in Supplementary Figure S4A. Therefore, the final temperature (immediately after the T-jump) was determined by finding $I_D(T_f)$ on the equilibrium profile such that the ratio $\frac{I_D(T_f)}{I_D(T_i)}$ matched the ratio $\frac{I(0^+)}{I(0^-)}$ obtained from the T-jump trace. The errors in the T-jump size are estimated to be about 10%.

The intensity $I(0^-)$ for each T-jump trace was obtained by averaging the pre-T-jump intensities prior to the arrival of the IR pulse. The intensity $I(0^+)$ was obtained by fitting the kinetics traces measured on donor-only samples to the so-called "T-jump recovery function":

$$I(t) = [I(0^+) - I(0^-)]f_{rec}(t) + I(0^-) \qquad \text{(S1)}$$

with $f_{rec}(t) = (1 + t/\tau_{rec})^{-1}$, where $\tau_{rec}$ is a characteristic time constant for the temperature of the heated volume of the sample to decay back to the initial equilibrium temperature (10). The recovery time constant $\tau_{rec}$ was determined for each sample as the average of several control measurements and found to be in the range of $102 - 210$ ms with an average value of $138 \pm 30$ ms.

**1.8 Single- or double-exponential decay coupled with T-jump recovery**. Relaxation kinetics traces $I(t)$ measured on donor-acceptor labeled samples were analyzed in terms of discrete exponential decays or using maximum entropy analysis, as described below. Discrete exponential analysis utilized either a single-exponential decay (with relaxation rate $k_r$; Eq. S2) or two-exponential decay (with relaxation rates $k_{fast}$ and $k_{slow}$; Eq. S3), combined with the recovery of the donor intensity back to the pre-T-jump levels characteristic of the initial temperature:

$$I(t) = \left(I(0^+) - I_{app}(\infty)\right) \times \exp(-k_r t) + \left(I_{app}(\infty) - I(0^-)\right) f_{rec}(t) + I(0^-) \qquad \text{(S2)}$$

$$I(t) = \left[I(0^+) - I_{app}(\infty)\right]\left[f_1 \exp\left(-k_{fast} t\right) + (1 - f_1)\exp\left(-k_{slow}(t)\right]\right.$$
$$+\left[I_{app}(\infty) - I(0^-)\right]f_{rec}(t) + I(0^-) \qquad \text{(S3)}$$

In the above equations, $I(0^-)$ and $I(0^+)$ are as defined above, $I_{app}(\infty)$ is the intensity at the end of the observed conformational relaxation process, and $f_1$ in Eq. S3 is the fractional amplitude in the fast component. The parameters that were varied in the fit are $I(0^+)$, $I_{app}(\infty)$, $k_r$ and $\tau_{rec}$ in Eq. S2 and $I(0^+)$, $I_{app}(\infty)$, $k_{fast}$, $k_{slow}$, $f_1$ and $\tau_{rec}$ in Eq. S3.

The two-exponential decay plus T-jump recovery function (Eq. S3) was also used for the initial matching and combining of the traces acquired over the different timescales. In this case, there were two additional parameters (multiplicative scale factors) that were applied to two of the traces to account for any systematic difference in the measured intensities of the three traces. Once appropriately scaled, the three data sets were combined into a single kinetic trace that was then used for all subsequent analyses.

**1.9 Maximum entropy method (MEM) for analyzing T-jump relaxation rates.** Model-independent distributions of relaxation time constants, $f(\log\tau)$, were inferred from the relaxation traces using the maximum entropy method (MEM) as implemented in the MemExp program (11,12). The $f$ distribution is obtained by maximizing its entropy $S$ while constraining the normalized residual sum of squares to be close to one (11,13,14). Logarithmically averaged and matched T-jump kinetics traces were input to the MEM analysis.

The entropy function was defined as (15):

$$S(f,F) = \sum_{j=1}^{M} \left[ f_j - F_j - f_j \ln(f_j / F_j) \right] \qquad \text{(S4)}$$

where $f_j$ are the discretized values of the distribution $f(\log\tau)$ and $F$ is the model distribution defaulted to in the absence of data; here, it was assumed to be a uniform, flat distribution.

MEM analysis on T-jump traces of all control samples yielded a single-peaked distribution (Supplementary Figure S4C). The average relaxation time for each distribution was computed as

$$< \log\tau > = \frac{\sum_{j=1}^{M} \log\tau_j \, f(\log\tau_j) d\log\tau_j}{\sum_{j=1}^{M} f(\log\tau_j) d\log\tau_j} \qquad \text{(S5)}$$

to obtain the characteristic recovery time constant $\tau_{rec} = 10^{<\log\tau>}$. These values were in the range of $\sim 108 - 269$ ms, consistent with those obtained from fits using Eq. S1.

For donor-acceptor-labeled samples that exhibited conformational relaxation in the T-jump window, the measured kinetics were concurrent with recovery. In most cases, the conformational kinetics and recovery components had opposite amplitudes corresponding to the scenario in which the post-T-jump intensities exhibited first a rising behavior, from $I(0^+)$ to $I_{app}(\infty)$, corresponding to conformational relaxation, and then a decaying behavior, from $I_{app}(\infty)$ to $I(0^-)$, corresponding to the T-jump recovery (Supplementary Figure S6A,C). For these cases, the distribution $f(\log\tau)$ was expressed as the sum of two contributions: $f(\log\tau) = g(\log\tau) - h(\log\tau)$, with $h(\log\tau)$ corresponding to the increase in intensity from $I(0^+)$ to $I_{app}(\infty)$ and $g(\log\tau)$ corresponding to the decrease in intensity from $I_{app}(\infty)$ to $I(0^-)$. The measured intensities were fitted to the following function form:

$$I(t) = D_o \int_{-\infty}^{+\infty} d\log\tau \, [g(\log\tau) - h(\log\tau)] \exp(-t/\tau) + I(0^-) \quad \text{(S6a)}$$

or the discretized version:

$$I(t) = D_0 \left( \sum_i [g(\log\tau_i) - h(\log\tau_i)] \, e^{-t/\tau_i} \right) d\log\tau + I(0^-) \quad \text{(S6b)}$$

In the above expression, $I(0^-)$ is the pre-T-jump intensity that was fixed from the data, $D_0$ is a normalization constant that was varied to best match the post-T-jump intensity level $I(0^+)$, and $d\log\tau$ is the separation between successive $\log\tau_i$ in the discrete MEM distribution.

The two different contributions to the MEM distribution for a given T-jump trace measured on a donor-acceptor sample are shown in Supplementary Figure S6B, together with the MEM distribution from a control experiment (on donor-only sample) that exhibits only the recovery kinetics. We note here that $h(\log\tau)$, which we assign as the distribution representing the conformational relaxation, and $g(\log\tau)$, which we assign as that for the T-jump recovery, appear cleanly separated in the MEM analysis. However, when the MEM is used to recover two lifetime distributions of opposite sign, its bias toward minimal structure (maximally uniform distributions) yields distributions of opposite sign that overlap minimally. A comparison between $g(\log\tau)$ obtained from a T-jump trace that has both contributions (black curve in Supplementary Fig. S6B) and that obtained from a control sample (gray curve in Supplementary Fig. S6B) shows that the latter peaks at shorter times and has a tail that extends into and overlaps with $h(\log\tau)$ (maroon curve in Supplementary Fig. S6B), demonstrating that relaxation and recovery, in fact, occur simultaneously. The current analysis, which ignores the tail at long times in the $h(\log\tau)$ distribution, therefore slightly overestimates the slow missing amplitudes. A more accurate treatment that decouples the two concurrent processes is beyond the scope of this work.

The two separate contributions to the measured T-jump trace are visualized in Supplementary Figure S6A (as computed from the discrete exponential fits described in Eqs. S2 or S3) and in Supplementary Figure S6C (as computed from the MEM outputs $h(\log\tau)$ and $g(\log\tau)$). For the MEM analysis, the conformational relaxation contribution ($I_{conf\_relaxation}$) is recapitulated as

$$I_{conf\_relaxation} = D_o \left(\sum_i g(\log\tau_i) - \sum_i h(\log\tau_i). e^{-t/\tau_i}\right) d\log\tau + I(0^-) \quad \text{(S7)}$$

and the T-jump recovery contribution ($I_{recovery}$) as

$$I_{recovery} = D_o \left(\sum_i g(\log\tau_i) e^{-t/\tau_i}\right) d\log\tau + I(0^-) \quad \text{(S8)}$$

The intensity levels at the beginning and end of the observed conformational relaxation, $I(0^+)$ and $I_{app}(\infty)$, respectively, were computed as

$$I(0^+) = I_{conf\_relaxation}(t \to 0) = D_o \sum_i [g(\log\tau_i) - h(\log\tau_i)]\, dlog\tau + I(0^-) \quad \text{(S9a)}$$

and

$$I_{app}(\infty) = I_{recovery}\,(t \to 0) = D_o \sum_i g(\log\tau_i)\, d\log\tau + I(0^-) \quad \text{(S9b)}$$

These intensity levels were further used for amplitude analysis, as described in Section 1.10.

For samples that exhibited the same sign for the relaxation kinetics and the recovery, i.e., the post-T-jump intensities increased from $I(0^+)$ to $I_{app}(\infty)$ and then further increased from $I_{app}(\infty)$ to $I(0^-)$ (Supplementary Figure S7A,C), the MEM distribution for the entire kinetic trace had amplitudes of the same sign, as illustrated in Supplementary Figure S7B. For these cases, the distributions obtained from MEM could still be separated into one distribution $h(\log\tau)$ corresponding to conformation relaxation and one distribution $g(\log\tau)$ corresponding to T-jump recovery, since the recovery contribution appeared as a distinct peak in the full MEM distribution and was readily identified by comparison with the MEM distribution of a donor-only control trace (Supplementary Figure S7B). In this case, the distribution corresponding to $h(\log\tau)$ was extracted from the full MEM distribution by selecting the left flank of the component from conformational relaxation, which was then mirrored to obtain a symmetric (in $\log\tau$ space)

$h(\log\tau)$ distribution. The recovery only component $g(\log\tau)$ was obtained by subtracting $h(\log\tau)$ as obtained above from the full MEM distribution. The time dependencies of $I_{conf\_relaxation}$ and $I_{recovery}$, and the intensities $I(0^+)$ and $I_{app}(\infty)$, were obtained from equations analogous to Eqs. S7-S9.

**1.10 Amplitude analysis.** The control (DNA_D) samples exhibit a decrease in donor quantum yield as a function of temperature, which is reflected in the temperature dependence of $I_D$ (Figure 3A and Supplementary Figure S5A; black symbols). For donor-acceptor (DNA_DA) samples, we anticipate a similar decrease in donor quantum yield when the temperature of the sample changes from $T_i$ to $T_f$, since the temperature response of the donor quantum yield is expected to be independent of whether there is an acceptor in the sample or not. Therefore, we expect the donor intensity in DNA_DA samples to decrease from $I_{DA}(T_i)$ to $\beta I_{DA}(T_i)$, where $\beta = \frac{I_D(T_f)}{I_D(T_i)}$, within the time that it takes for the sample to heat up after the arrival of the IR pulse (Figure 3A and Supplementary Figure S5A; dashed red line, computed as $I_D(T) \times \frac{I_{DA}(T_i)}{I_D(T_i)}$). This drop is illustrated by the black arrow on the equilibrium profile shown in Figure 3A. Next, the donor intensity level is expected to reach that of $I_{DA}(T_f)$ because of DNA conformational changes (illustrated by the purple arrow in Figure 3A Supplementary Figure S5A), if the conformational relaxation reaches completion before the T-jump itself starts to recover. Finally, the intensity levels are expected to decay back to $I_{DA}(T_i)$ when the temperature decays back. Therefore, the maximum amplitude we expect for the conformational relaxation kinetics in the T-jump traces can be computed as $Amp_{eq} = \frac{I_{DA}(T_f) - \beta I_{DA}(T_i)}{I_{DA}(T_i)}$, where we have normalized the amplitude to the initial intensity level $I_{DA}(T_i)$.

This estimation of total expected amplitude is illustrated in Supplementary Figure S5A for TAT/TAT DNA, for a T-jump from $T_i = 13$ ℃ to $T_f = 19$ ℃. Here we show the intensities normalized to that at the initial temperature: $I_{DA}^N = \frac{I_{DA}}{I_{DA}(T_i)}$ and $I_D^N = \frac{I_D}{I_D(T_i)}$. In terms of these normalized intensities, the total expected amplitude becomes $Amp_{eq} = I_{DA}^N(T_f) - \beta = I_{DA}^N(T_f) - I_D^N(T_f)$, illustrated by the purple arrow. A larger T-jump size would lead to a larger expected amplitude. The amplitudes thus calculated for two different T-jump sizes (6 ℃ and 9 ℃) on the TAT/TAT sample are shown in Supplementary Figure S5C as a function of the final temperature $T_f$. The data point corresponding to the T-jump conditions of Supplementary Figure S5A is circled in purple.

To account for variations in expected amplitudes from variations in the T-jump size from one set of measurements to another, we divide the computed amplitudes by a scale factor $\frac{I_D(T_i)-I_D(T_f)}{I_D(T_i)} = 1 - I_D^N(T_f)$, which is equal to the length of the black arrow in Figure 3A and Supplementary Figure S5A. Since $I_D$ decreases approximately monotonically with temperature, this scale factor is roughly proportional to the size of the T-jump. With this correction, we obtain the appropriately scaled (for the size of the T-jump) amplitudes: $Amp_{eq}^s = \frac{Amp_{eq}}{1-I_D^N(T_f)}$, plotted in Supplementary Figure S5E. The circled data point has a scaled amplitude of ~3, consistent with the equilibrium profiles in Supplementary Figure S5A, where the purple arrow (change in intensity expected from conformational dynamics for a given T-jump size) is ~3-fold larger than the black arrow (change in intensity from donor quantum yield changes for that T-jump).

The donor intensity levels measured in the equilibrium steady-state measurements can be mapped over to the donor intensity levels we expect to measure in our T-jump spectrometer. For example, immediately after the T-jump and prior to any conformational dynamics, we expect the intensity levels to jump from $I(0^-)$, the pre-T-jump intensity level, to $I_{eq}(0^+)$, indicated by the black dashed line in Figure 3B and Supplementary Figure S5B), and computed as $I_{eq}(0^+) = I(0^-) \times \beta = I(0^-) \times I_D^N(T_f)$, since $I_D^N(T_f)$ is a measure of the fractional change in the donor quantum yield as a result of the T-jump. Similarly, the donor intensity level we expect at the end of the conformational relaxation, assuming much slower T-jump recovery (denoted as $I_{eq}(\infty)$; red dashed line in Figure 3B and Supplementary Figure S5B), was computed as $I_{eq}(\infty) = I(0^-) \times \frac{I_{DA}(T_f)}{I_{DA}(T_i)} = I(0^-) \times I_{DA}^N(T_f)$, where $I_{DA}^N(T_f)$ is a measure of the fractional change in the donor intensity of the DNA_DA samples as a result of the change in the equilibrium populations from the initial temperature distribution to that at the final temperature. Defined in terms of the intensities measured in the T-jump spectrometer, the total expected amplitude is equivalently written as $Amp_{eq} = \frac{I_{eq}(\infty)-I_{eq}(0^+)}{I(0^-)}$, which is equal to the vertical purple arrow in Figure 3B and Supplementary Figure S5B.

However, not all the amplitude expected from equilibrium measurements is in fact observed within our T-jump time window, as illustrated in Figure 3B and Supplementary Figure S5B. In the T-jump trace shown for a donor-acceptor-labeled (DNA_DA) sample, the observed donor intensity immediately after the T-jump (denoted as $I(0^+)$; green dashed line in Figure 3B and Supplementary Figure S5B) does not drop as low as we expected, i.e., the post-T-jump intensity does not reach the level of $I_{eq}(0^+)$ as defined above. This difference between $I_{eq}(0^+)$ and $I(0^+)$ (vertical maroon arrow in Figure 3B and Supplementary Figure S5B) is assumed to be from an unresolved "missing" fast phase, with amplitude $Amp_{fast} = \frac{I(0^+) - I_{eq}(0^+)}{I(0^-)}$, where again we have normalized the amplitude to the pre-T-jump level $I(0^-)$.

Similarly, the intensity level we measure at the apparent end of the observed relaxation phase (denoted as $I_{app}(\infty)$; cyan dashed line in Figure 3B and Supplementary Figure S5B) does not reach the anticipated level $I_{eq}(\infty)$, presumably because the temperature of the sample starts to recover before the completion of the conformational relaxation. In this case, the difference between $I_{eq}(\infty)$ and $I_{app}(\infty)$ (vertical cyan arrow in Figure 3B and Supplementary Figure S5B) is defined as the amplitude of the "missing" slow phase, with amplitude $Amp_{slow} = \frac{I_{eq}(\infty) - I_{app}(\infty)}{I(0^-)}$. The observed amplitude (vertical green arrow) is then $Amp_{obs} = \frac{I_{app}(\infty) - I(0^+)}{I(0^-)}$ such that $Amp_{fast} + Amp_{obs} + Amp_{slow} = Amp_{eq}$ adds up to the total amplitude expected from the equilibrium measurements.

The calculation of $Amp_{obs}$ from the relaxation trace measured for the TAT/TAT sample, for a T-jump from $T_i = 13$ ℃ to $T_f = 19$ ℃, is illustrated in Supplementary Figure S5B. The $I(0^+)$ and $I_{app}(\infty)$ levels were computed as described in SI Methods 1.9. The intensities plotted here are normalized such that $I(0^-) = 1$ . In this case, the observed amplitude $Amp_{obs}$ is simply the length of the vertical green arrow. The observed amplitudes for the TAT/TAT sample for two sets of measurements are shown in Supplementary Figure S5D, with the data point corresponding to the shown relaxation trace encircled in green. Again, to account for variations in the T-jump size, the observed amplitudes (as well as the fast and slow "missing" amplitudes) were normalized by the T-jump scale factor $1 - I_D^N(T_f)$ to obtain the scaled amplitudes $Amp_{obs}^s$, $Amp_{fast}^s$, and $Amp_{slow}^s$, which as discussed above is equivalent to dividing the magnitudes of the measured amplitudes: green (observed), maroon (fast), and cyan (slow) in Figure 3 – by the total change in the quantum yield of the donor (black arrow). The appropriately scaled amplitudes observed in the T-jump traces of the TAT/TAT samples are shown in Supplementary Figure S5F.

The amplitudes presented in the main text are the average of two sets of measurements on each sample. The differences in T-jump size from one set of measurements to the next translate into differences in the final temperature between the two sets. Therefore, prior to averaging the amplitudes, they were interpolated to a common (final) temperature grid, from 20 ℃ to 40 ℃ in steps of 5 ℃, as illustrated in Supplementary Figure S5G-H. The interpolated amplitudes were then averaged to obtain the data shown in Supplementary Figure S5I-J and correspond to the amplitudes shown in the main text.

In rare cases, primarily at the low temperature end, the observed amplitude in the T-jump trace ($Amp_{obs}$) was greater than the expected amplitude from the equilibrium data ($Amp_{eq}$); see for example the 20 ℃ data in Figure 4C & 4I. This situation arose if $I(0^+)$ fell below $I_{eq}(0^+)$ (see Figure S9F) or if $I_{app}(\infty)$ exceeded $I_{eq}(\infty)$ (see Figure S8F and Figure S9F), leading to apparently negative amplitudes for the missing fast or missing slow phases. This unphysical scenario was attributed to distortions in the equilibrium profiles at low temperatures, perhaps from some condensation effects in the sample chamber of the steady-state fluorescence spectrometer, leading to systematic errors in the estimation of the T-jump size and/or the estimation of the computed $I_{eq}(0^+)$ and $I_{eq}(\infty)$ levels. The negative amplitudes were zeroed out by forcing the $I_{eq}(0^+)$ levels to be equal to the observed $I(0^+)$ values or the $I_{eq}(\infty)$ levels to be equal to $I_{app}(\infty)$ values and recalculating the amplitudes.

**1.11 Error propagation in the computed amplitudes**. A major source of error in the expected amplitudes computed from the equilibrium thermal profiles is the uncertainty in the measurements of the thermal profiles $I_D$ and $I_{DA}$. The uncertainties in the raw measured intensities (denoted as $\sigma_I$ and computed as standard error of the mean, s.e.m, from independent runs on a given sample) are shown in Supplementary Figure S2B. To account for variations in the measured intensities from one day to the next, all measured intensities, $I_D$ and $I_{DA}$, were normalized with respect to the intensity $I_{DA}$ at 10 ℃ on that day prior to computing the averages and s.e.m values. In this case, the uncertainties in $I_{DA}$ at 10 ℃ are, by definition, zero. For the thermal profiles $I_D^M$ and $I_{DA}^M$ shown in Figure 2 and Supplementary Figure S2C, the intensities $I_D$

from independent runs were also first normalized with respect to the intensity $I_D$ at 10 ℃ such that $I_D^M = I_{DA}^M = 1$ at 10 ℃; the uncertainties at temperatures greater than 10 ℃ (denoted as $\sigma_I^M$) were calculated from the s.e.m. of the data matched in this way.

For amplitude analysis, what is relevant is the uncertainties in $I_D(T_f)$ and $I_{DA}(T_f)$ relative to the corresponding intensities at the initial temperature, $I_D(T_i)$ and $I_{DA}(T_i)$, respectively. In this case, the measured thermal profiles were normalized to match at the initial temperature, denoted as $I_D^N$ and $I_{DA}^N$ with corresponding s.e.m. $\sigma_I^N$ (equal to zero at $T_i$ by definition), as shown in Supplementary Figure S5A. These errors, from measurements on DNA_D and DNA_DA samples, appeared in the calculation of the errors in the expected intensity levels $I_{eq}(0^+)$ and $I_{eq}(\infty)$, respectively, for a given set of T-jump conditions, and propagated into the calculation of the errors in the expected total amplitude, as shown in Supplementary Figure S5C, as well as into the calculation of the errors in the missing fast and slow amplitudes. Each data set, after correcting for the T-jump size (illustrated in Supplementary Figure S5E) and interpolated on a common temperature grid (illustrated in Supplementary Figure S5G) then has an associated error in the calculated amplitudes. For a given data set $i$ with computed amplitude $A_i$ and uncertainty $\sigma_{Ai}$, the weighted average $A'$ and the weighted variance of the mean $\sigma_{A'}^2$ (from $N$ independent runs) were calculated as $A' = \frac{\sum A_i/\sigma_{Ai}^2}{\sum 1/\sigma_{Ai}^2}$ and $\sigma_{A'}^2 = \frac{1}{(N-1)} \frac{\sum (A_i - A')^2/\sigma_{Ai}^2}{\sum 1/\sigma_{Ai}^2}$. Finally, to include the contribution of errors from both the variance of the mean (dominated by statistical fluctuations from one data set to another) and errors in the amplitudes in each data set $i$ (dominated by errors in the thermal melting profiles), the two sources of errors were combined as $\sigma_A^2 = \sigma_{A'}^2 + \sum \sigma_{Ai}^2/N$; these are the errors plotted in Supplementary Figure S5I).

For computation of errors in the observed amplitudes, the errors in the amplitudes from a given T-jump trace were assumed to be negligible in comparison with the variability from one data set to another (see Supplementary Figure S5D, F, H). Therefore, the mean and s.e.m. for the observed amplitudes were simply the corresponding unweighted quantities from 2 independent runs (illustrated in Supplementary Figure S5J).

### SI References


1. Cantor, C.R., Warshaw, M.M. and Shapiro, H. (1970) Oligonucleotide interactions. 3. Circular dichroism studies of the conformation of deoxyoligonucleotides. *Biopolymers*, **9**, 1059-1077.
2. Cavaluzzi, M.J. and Borer, P.N. (2004) Revised UV extinction coefficients for nucleoside-5'-monophosphates and unpaired DNA and RNA. *Nucleic Acids Res*, **32**, e13.
3. Min, J.H. and Pavletich, N.P. (2007) Recognition of DNA damage by the Rad4 nucleotide excision repair protein. *Nature*, **449**, 570-575.
4. Chen, X., Velmurugu, Y., Zheng, G., Park, B., Shim, Y., Kim, Y., Liu, L., Van Houten, B., He, C., Ansari, A. *et al.* (2015) Kinetic gating mechanism of DNA damage recognition by Rad4/XPC. *Nat Commun*, **6**, 5849.
5. Clegg, R.M. (1992) Fluorescence resonance energy transfer and nucleic acids. *Methods Enzymol.*, **211**, 353-388.
6. Preus, S., Borjesson, K., Kilsa, K., Albinsson, B. and Wilhelmsson, L.M. (2010) Characterization of nucleobase analogue FRET acceptor tCnitro. *J Phys Chem B*, **114**, 1050-1056.
7. Chakraborty, S., Steinbach, P.J., Paul, D., Mu, H., Broyde, S., Min, J.H. and Ansari, A. (2018) Enhanced spontaneous DNA twisting/bending fluctuations unveiled by fluorescence lifetime distributions promote mismatch recognition by the Rad4 nucleotide excision repair complex. *Nucleic Acids Res*, **46**, 1240-1255.
8. Velmurugu, Y., Chen, X., Slogoff Sevilla, P., Min, J.H. and Ansari, A. (2016) Twist-open mechanism of DNA damage recognition by the Rad4/XPC nucleotide excision repair complex. *Proc Natl Acad Sci U S A*, **113**, E2296-2305.
9. Wray, W.O., Aida, T. and Dyer, R.B. (2002) Photoacoustic cavitation and heat transfer effects in the laser-induced temperature jump in water. *Appl. Phys. B*, **74**, 57-66.
10. Vivas, P., Kuznetsov, S.V. and Ansari, A. (2008) New insights into the transition pathway from nonspecific to specific complex of DNA with Escherichia coli integration host factor. *J Phys Chem B*, **112**, 5997-6007.
11. Steinbach, P.J., Ionescu, R. and Matthews, C.R. (2002) Analysis of kinetics using a hybrid maximum-entropy/nonlinear-least-squares method: application to protein folding. *Biophys. J.*, **82**, 2244-2255.

12. Sternisha, S.M., Whittington, A.C., Martinez Fiesco, J.A., Porter, C., McCray, M.M., Logan, T., Olivieri, C., Veglia, G., Steinbach, P.J. and Miller, B.G. (2020) Nanosecond-Timescale Dynamics and Conformational Heterogeneity in Human GCK Regulation and Disease. *Biophys J*, **118**, 1109-1118.
13. Livesey, A.K. and Brochon, J.C. (1987) Analyzing the Distribution of Decay Constants in Pulse-Fluorimetry Using the Maximum Entropy Method. *Biophys J*, **52**, 693-706.
14. Steinbach, P.J. (2002) Inferring lifetime distributions from kinetics by maximizing entropy using a bootstrapped model. *J Chem Inf Comput Sci*, **42**, 1476-1478.
15. Skilling, J. (1989) In Skilling, J. (ed.), *Maximum Entropy and Bayesian Methods*. Springer Dordrecht, pp. 45-52.
16. Paul, D., Mu, H., Zhao, H., Ouerfelli, O., Jeffrey, P.D., Broyde, S. and Min, J.H. (2019) Structure and mechanism of pyrimidine-pyrimidone (6-4) photoproduct recognition by the Rad4/XPC nucleotide excision repair complex. *Nucleic Acids Res*, **47**, 6015-6028.

5′-TTGACTCGACATCPXXXGGTACAA-3′
3′-ACTGAGCTGTAGGYYYCDATGTTA-5′

| Sample<br>XXX/YYY | $K_{d,app}$ (nM) | $T_m$ (°C) | FRET E (10 °C) | FRET E (40 °C) |
|---|---|---|---|---|
| CCC/CCC | 70 ± 3 | 60.6 ± 1.8 | 0.81 ± 0.01<br>(0.63 ± 0.01) | 0.73 ± 0.01<br>(0.54 ± 0.001) |
| GGG/CCC | 355 ± 28 | 78.6 ± 1.0 | 0.93 ± 0.01 | 0.92 ± 0.01 |
| TTT/TTT | 116 ± 3 | 62.1 ± 0.5 | 0.89 ± 0.01<br>(0.85 ± 0.01) | 0.80 ± 0.01<br>(0.75 ± 0.01) |
| AAA/TTT | 377 ± 9 | 72.7 ± 1.1 | 0.91 ± 0.01 | 0.87 ± 0.01 |
| TAT/TAT | 504 ± 36 | 65.6 ± 0.9 | 0.86 ± 0.01<br>(0.86 ± 0.01) | 0.79 ± 0.01<br>(0.78 ± 0.01) |
| ATA/TAT | 534 ± 44 | 74.4 ± 0.4 | 0.92 ± 0.01 | 0.87 ± 0.01 |
| 6-4 PP | 35 ± 1 | 61.30 | - | - |
| CPD | 302 ± 26 | 69.20 | - | - |

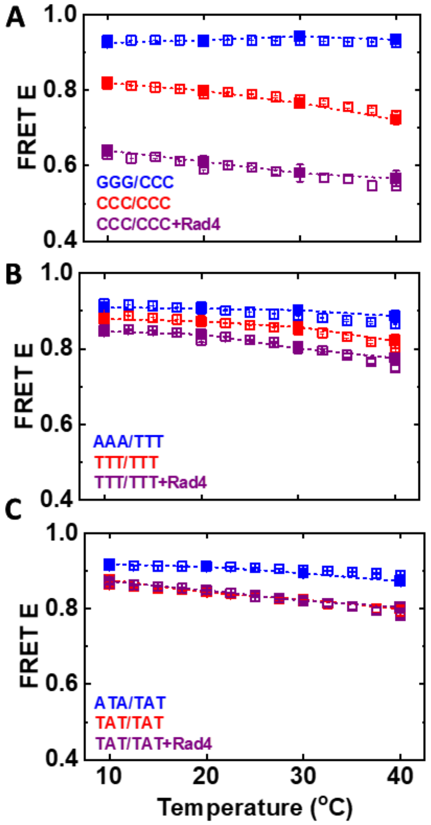


**Figure S1. Binding affinities, melting temperatures, and FRET measurements on tC°-tC$_{nitro}$ labeled DNA.** The DNA sequence used in this study is shown with the locations of the donor tC° (D in blue) and the acceptor tC$_{nitro}$ (P in cyan) indicated. The XXX/YYY in the sequence is replaced by CCC/CCC, GGG/CCC, etc, to create the different mismatched/matched constructs listed in the Table. The table summarizes the apparent binding affinities ($K_{d,app}$) of Rad4 for each of these DNA constructs and their melting temperatures ($T_m$), from ref. (7), and for NER lesions 6-4 photoproduct (6-4 PP) and cyclobutane pyrimidine dimer (CPD), from ref. (16). The average FRET efficiencies (FRET E) obtained from steady-state fluorescence measurements are also tabulated for two different temperatures; the numbers in parenthesis are in the presence of Rad4. (A-C) The average FRET efficiencies measured using two different methods – steady-state measurements (open symbols) and fluorescence lifetime measurements (filled symbols) – are plotted as a function of temperature for (A) the CCC/CCC data set; (B) the TTT/TTT data set; and (C) the TAT/TAT data set. In each panel data are shown for matched DNA (blue), mismatched DNA (red), and mismatched+Rad4 (purple).

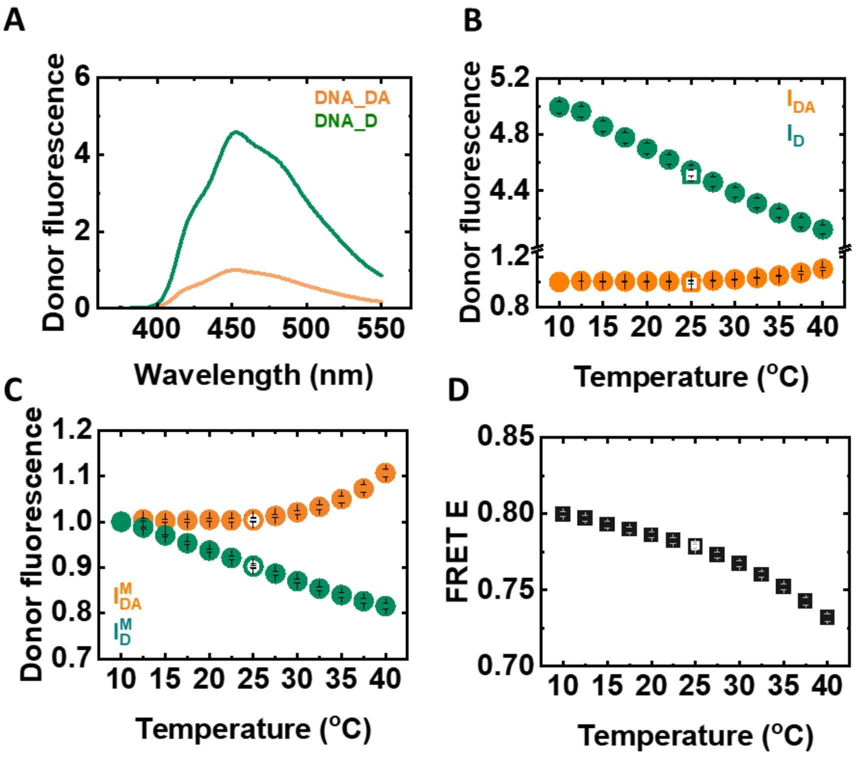


**Figure S2. Steady-state fluorescence measurements.** (A) Donor (tC°) fluorescence emission spectra are shown for the mismatched CCC/CCC construct at 25 ℃, measured in the absence (DNA_D; green) and presence (DNA_DA; orange) of the acceptor ($tC_{nitro}$). (B) Donor emission intensities, computed from the area under the emission spectra in the wavelength range 425 – 475 nm, are plotted as a function of temperature for donor-only ($I_D$; green) and donor-acceptor ($I_{DA}$; orange) samples. (C) The intensities $I_D$ and $I_{DA}$ from panel (B) were normalized to match at 10 ℃. (D) The FRET efficiencies, computed as described in SI Methods 1.3, are plotted as a function of temperature. Measurements were done from 10 ℃ up to 40 ℃ with 2.5 ℃ interval (filled symbols) and then the samples were cooled back to 25 ℃ and measured again (open symbols) to check for sample reversibility.

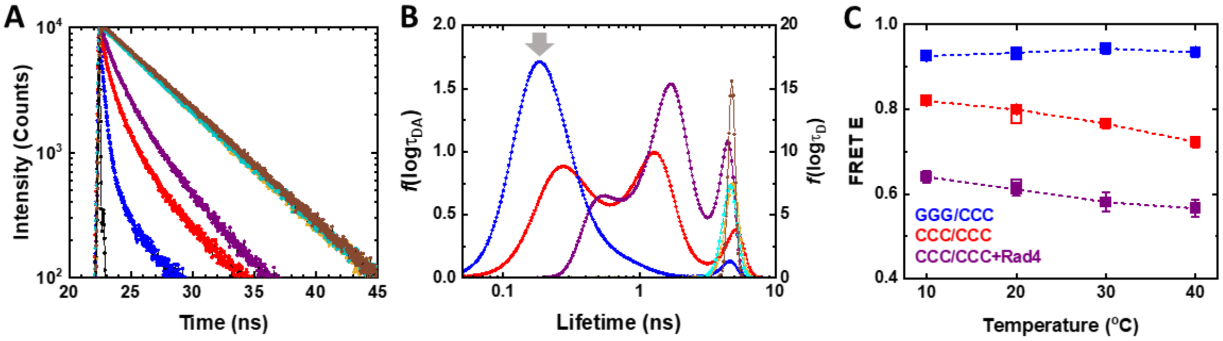


**Figure S3. Fluorescence lifetime measurements.** (A) Fluorescence intensity decay curves for DNA_DA (labeled with donor and acceptor) with excitation of donor (tC°) are shown for matched GGG/CCC (blue), mismatched CCC/CCC (red), and CCC/CCC+Rad4 (purple) constructs; the corresponding intensity decay curves for DNA_D (labeled only with donor) are shown in cyan, mustard, and brown respectively. The instrument response function (IRF) is shown in black. (B) The distribution of lifetimes that best describe the intensity decay profiles, as obtained from the maximum entropy method (MEM), are shown. The amplitudes from the MEM analysis, normalized to add up to one, are indicated on the left $y$-axis for DNA_DA and on the right $y$-axis for DNA_D. The gray arrow indicates the lifetime corresponding to a FRET of 0.92 expected for B-DNA conformations given the placements of the tC° and $tC_{nitro}$ FRET probes, as described in ref. (7). (C) The FRET efficiencies, computed as described in the text, are plotted as a function of temperature. Measurements were done from 10 ℃ up to 40 ℃ with 10 ℃ interval (filled symbols) and then the samples were cooled back to 20 ℃ and measured again (open symbols) to check for sample reversibility.

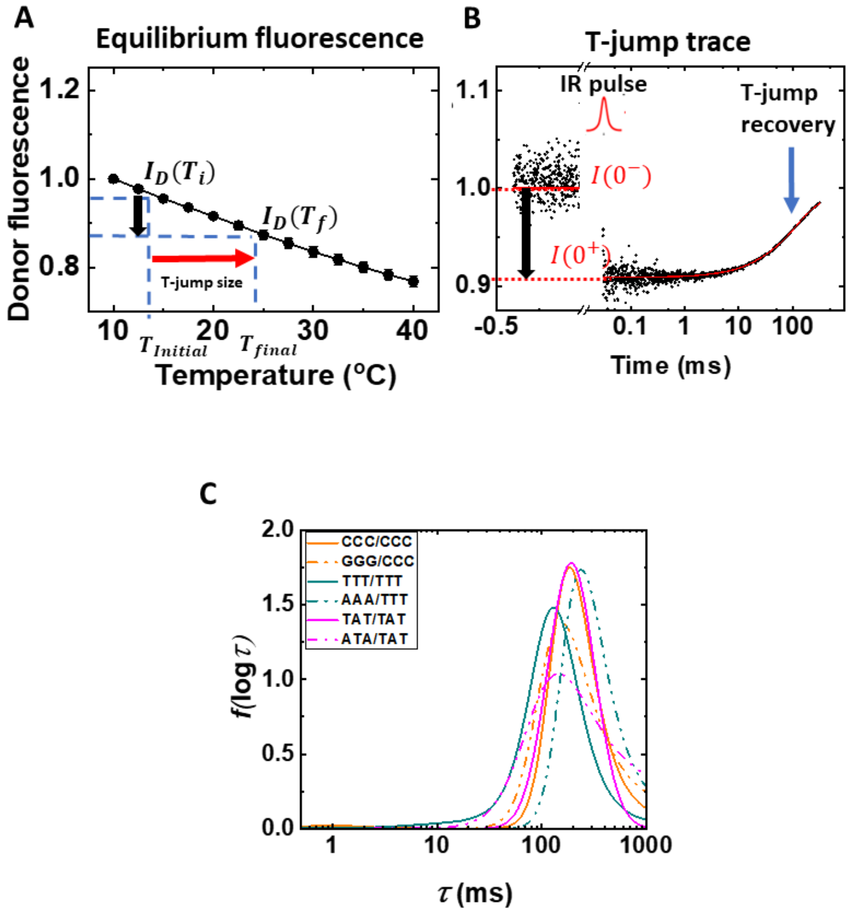


**Figure S4. Steady-state equilibrium and T-jump measurements on donor-only control samples.** (A) The fluorescence emission intensities $I_D$ measured at different temperatures are plotted as a function of temperature for all DNA_D constructs. The intensities are normalized to match at the lowest temperature. The black vertical arrow pointing down illustrates the decrease in donor intensity for donor-only (DNA_D) samples because of the decrease in the donor quantum yield when the temperature of the sample jumps from $T_{initial}$ $(T_i)$ to the final temperature $T_{final}$ $(T_f)$. (B) A typical T-jump trace for a donor-only DNA_D construct is shown. The donor intensity before the arrival of the IR pulse is at the pre-flash level $I(0^-)$. The donor intensity immediately after the T-jump is at a lower level $I(0^+)$ and this drop is attributed to the donor quantum yield change when the temperature of the sample jumps from $T_i$ to $T_f$. The donor intensity then decays back to $I(0^-)$ as the temperature of the sample decays back to $T_i$. The level $I(0^+)$ is determined by fitting the T-jump trace to the T-jump recovery function Eq. S5. The final temperature $T_f$ is determined by computing the intensity $I_D(T_f)$ such that $\frac{I_D(T_f)}{I_D(T_i)} = \frac{I(0^+)}{I(0^-)}$ and interpolating $I_D(T_f)$ on the $I_D$ versus temperature plot to find the corresponding temperature $T_f$. (C) The distribution of relaxation times obtained from MEM analysis of the T-jump traces on all the control (DNA_D) samples are shown.

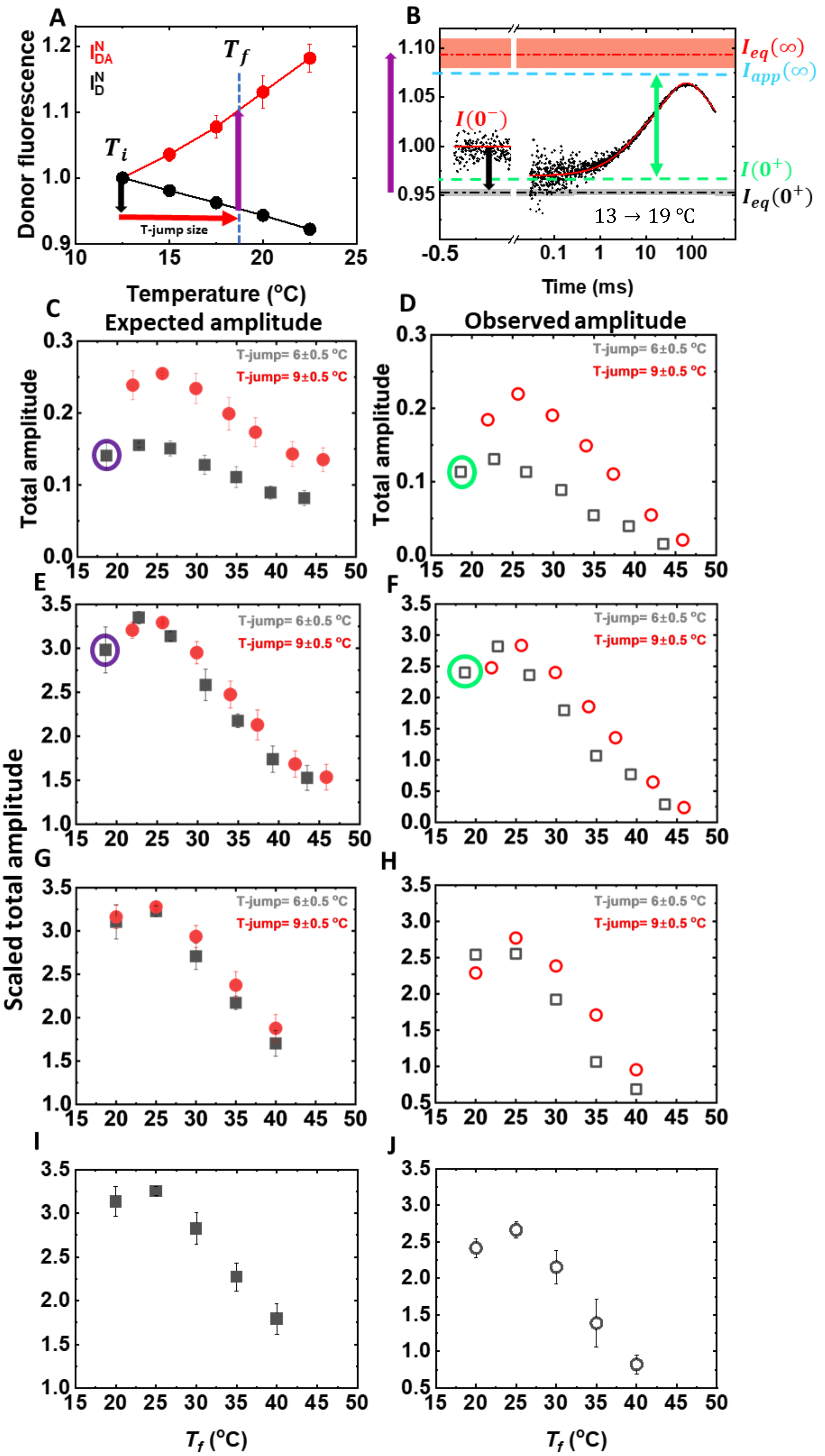

A
B
C
D
E
F
G
H
I
J
Donor fluorescence
Temperature (°C)
T-jump size
$T_i$
$T_f$
$I_{eq}(\infty)$
$I_{app}(\infty)$
$I(0^-)$
$I(0^+)$
$I_{eq}(0^+)$
13 → 19 °C
Time (ms)
Expected amplitude
Observed amplitude
Total amplitude
Scaled total amplitude
T-jump= 6±0.5 °C
T-jump= 9±0.5 °C
$T_f$ (°C)

**Figure S5. Illustration of amplitude analysis from steady-state equilibrium and T-jump measurements.** (A) The fluorescence emission intensities, after normalization to match at the initial temperature ($T_i$), are plotted as a function of temperature for DNA_D ($I_D^N$; black) and DNA_DA ($I_{DA}^N$; red). The black vertical arrow illustrates the drop in donor intensity in DNA_D samples when the temperature of the sample jumps from the initial ($T_i$) to the final ($T_f$) temperature. It is also the drop expected in DNA_DA samples immediately after the T-jump (prior to any conformational changes), as described in SI Methods 1.10. The blue vertical line indicates the intensity levels in the two samples under equilibrium conditions at $T_f$ relative to the intensity at $T_i$. (B) A representative T-jump trace is shown for a DNA_DA sample, with the relevant intensity levels indicated: $I(0^-)$ (continuous red), $I(0^+)$ (dashed green), $I_{app}(\infty)$(dashed cyan), $I_{eq}(0^+)$ (dashed black), and $I_{eq}(\infty)$ (dashed red). The uncertainties in $I_{eq}(0^+)$ and $I_{eq}(\infty)$, propagated from the equilibrium melting profiles as described in SI Methods 1.11, are shown as gray and red shaded regions, respectively. The vertical arrows indicate the total expected amplitude (purple) and the observed amplitude (green), computed as described in SI Methods 1.10. (C-D) The total expected (C) and observed (D) amplitudes for two different T-jump sizes for a given sample are plotted as a function of the final temperature. The errors in the expected amplitudes are computed from the errors in the equilibrium thermal profiles, as described in SI Methods 1.11. (E, F) The scaled amplitudes from two different sets of measurements, after correcting for variations in the T-jump size, are shown. (G, H) The amplitudes shown are from two different sets after interpolating onto a common temperature grid, prior to averaging. (I, J) The averaged amplitudes from two sets of measurements and the corresponding errors, computed as described in SI Methods 1.11, are shown.

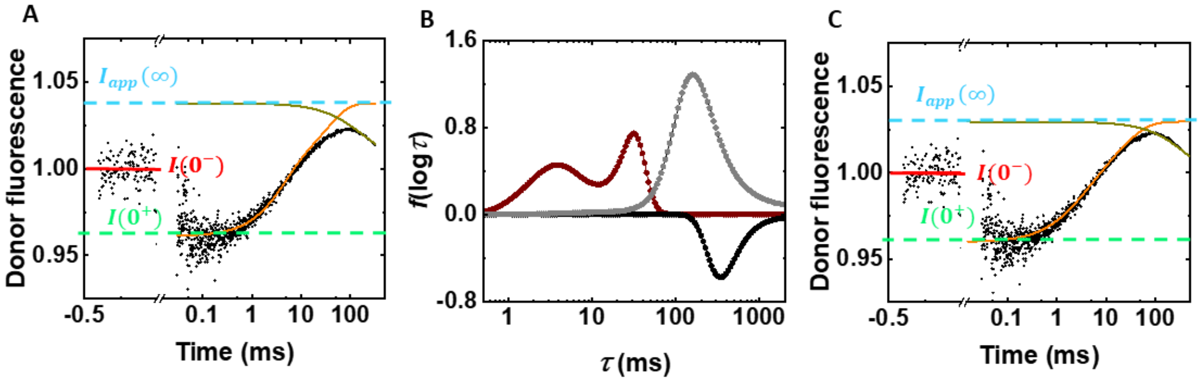


**Figure S6. T-jump traces where conformational relaxation and T-jump recovery have amplitudes of opposite signs.** (A) An example of a T-jump kinetic trace where the donor intensity measured after the arrival of the IR pulse starts at $I(0^+)$, then increases and overshoots past $I(0^-)$ to the intensity level $I_{app}(\infty)$, before decaying back to $I(0^-)$ as the temperature recovers. The black continuous line is a fit to the relaxation trace using the two-exponential decay function convoluted with the T-jump recovery (Eq. S3). The conformational relaxation contribution ($I_{conf\_relaxation}$; orange continuous line) and the T-jump recovery contribution ($I_{recovery}$; olive continuous line) are also shown separately. (B) The distribution of the logarithm of the relaxation times $f(\log\tau)$ obtained from MEM analysis of the relaxation trace is plotted; the positive amplitudes (maroon) describe the distribution $h(\log\tau)$ that corresponds to the conformational relaxation kinetics; the negative amplitudes (black) describe the distribution $g(\log\tau)$ that corresponds to the decay of the T-jump. The opposite signs for the two functions reflect the behavior of the kinetic trace in which the donor intensity first increases and then decreases. The gray curve is the MEM distribution obtained from a T-jump measurement on a control (DNA_D) sample. (C) The kinetic trace is as shown for panel (A); the continuous lines are computed from the MEM fits as described in the text: $I(t)$ from Eq. S6 (black); $I_{conf\_relaxation}$ from Eq. S7 (orange); $I_{recovery}$ from Eq. S8 (olive).

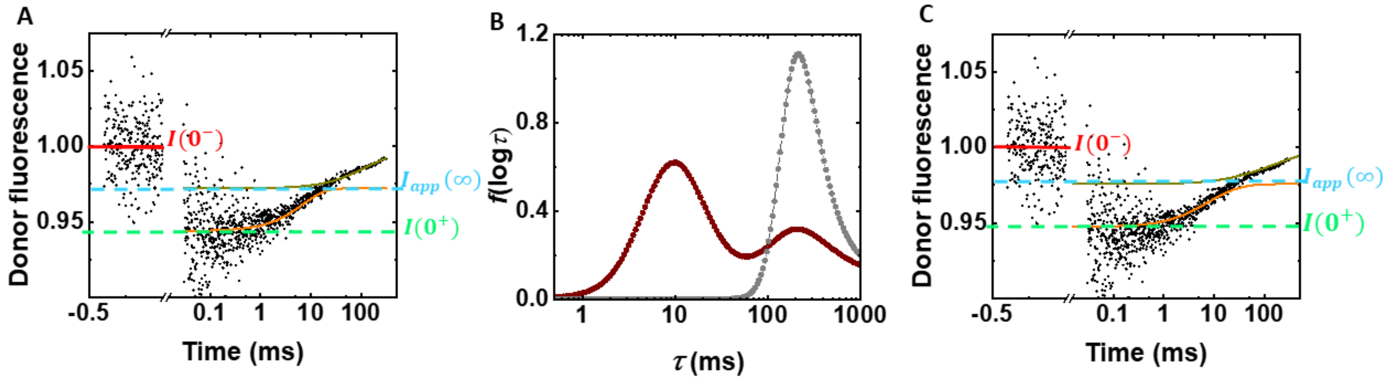


**Figure S7. T-jump traces where conformational relaxation and T-jump recovery have amplitudes of the same sign.** (A) An example of a T-jump kinetic trace where the donor intensity $I(0^+)$ measured immediately after the arrival of the IR pulse increases but does not overshoot past $I(0^-)$; instead, it reaches the intensity level $I_{app}(\infty)$ and then continues to increase to reach $I(0^-)$ as the temperature eventually decays back to the initial temperature. (B) The distribution $f(\log\tau)$ obtained from MEM analysis of the relaxation trace is plotted (maroon); all amplitudes, whether from conformational relaxation kinetics or from T-jump recovery, have the same sign. The two contributions $g(\log\tau)$ and $h(\log\tau)$ are separated as described in the text. The gray curve is the MEM distribution obtained from a T-jump measurement on a control (DNA_D) sample. (C) The kinetic trace is as shown for panel (A); the continuous lines are as described for Figure S6C.

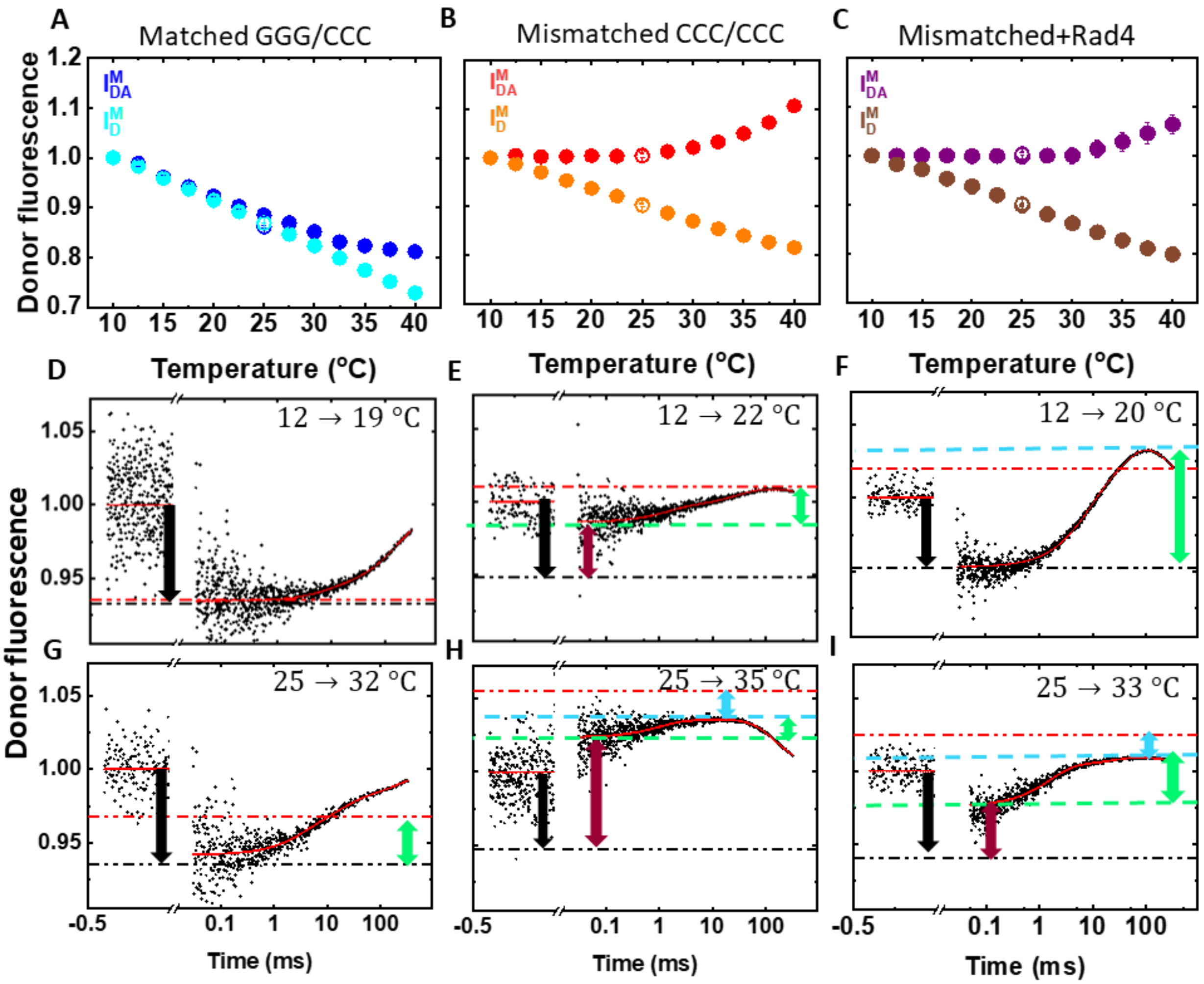


**Figure S8. Equilibrium and T-jump measurements on the CCC/CCC dataset. (A-C)** The donor fluorescence emission intensities $I_D/I_{DA}$ are plotted as a function of temperature, for (A) GGG/CCC (cyan/blue), (B) CCC/CCC (mustard/red), and (C) CCC/CCC+Rad4 (brown/purple). In each panel, the intensities $I_D$ and $I_{DA}$ have been normalized to match at 10 °C. (D-I) Donor fluorescence emission intensities of double-labeled (DNA_DA) samples, measured in response to a T-jump perturbation, are plotted as a function of time; final temperatures after the T-jump are in the range 19-22 °C (D-F) and 32-35 °C (G-I). In panels (D-I), the continuous red lines are from MEM fits to the relaxation traces; the horizontal lines and the vertical arrows are as described for Figure 3B and Supplementary Figure S5B.

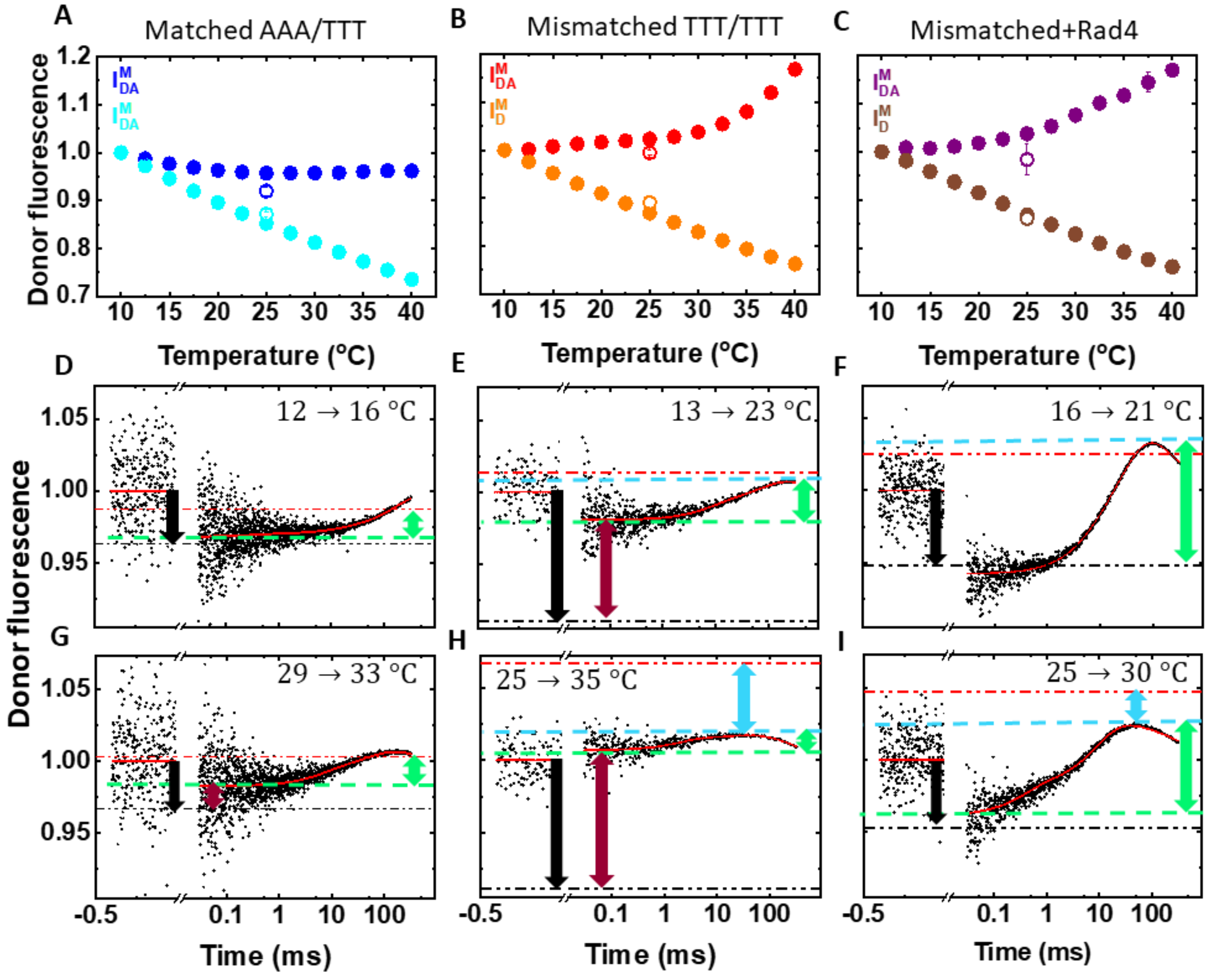


**Figure S9. Equilibrium and T-jump measurements on the TTT/TTT dataset.** Panels are as described for Figure S8. Final temperatures after the T-jump are in the range 16-23 °C (D-F) and 30-35 °C (G-I).

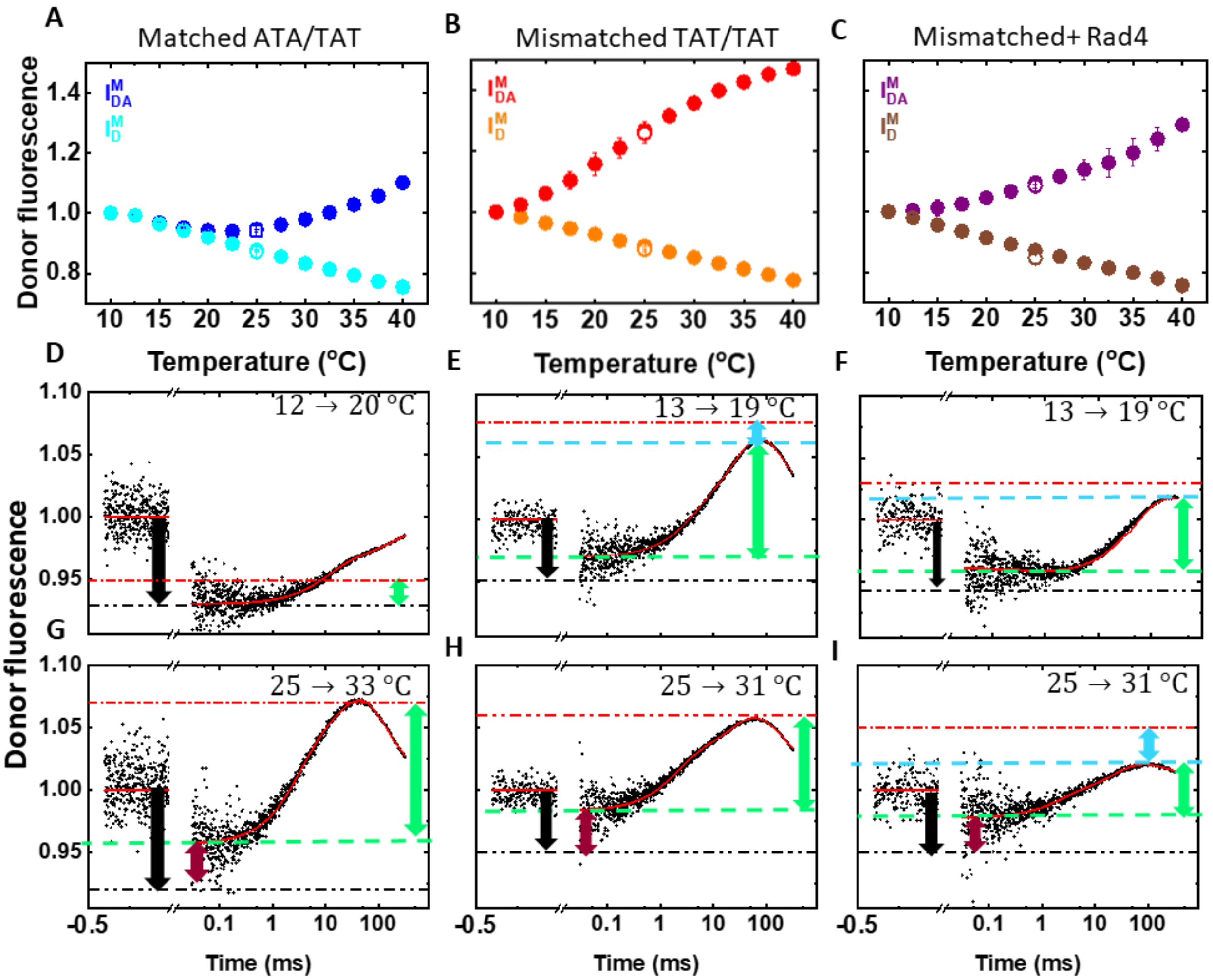


**Figure S10. Equilibrium and T-jump measurements on the TAT/TAT dataset.** Panels are as described for Figure S8. Final temperatures after the T-jump are in the range 19-20 °C (D-F) and 31-33 °C (G-I).

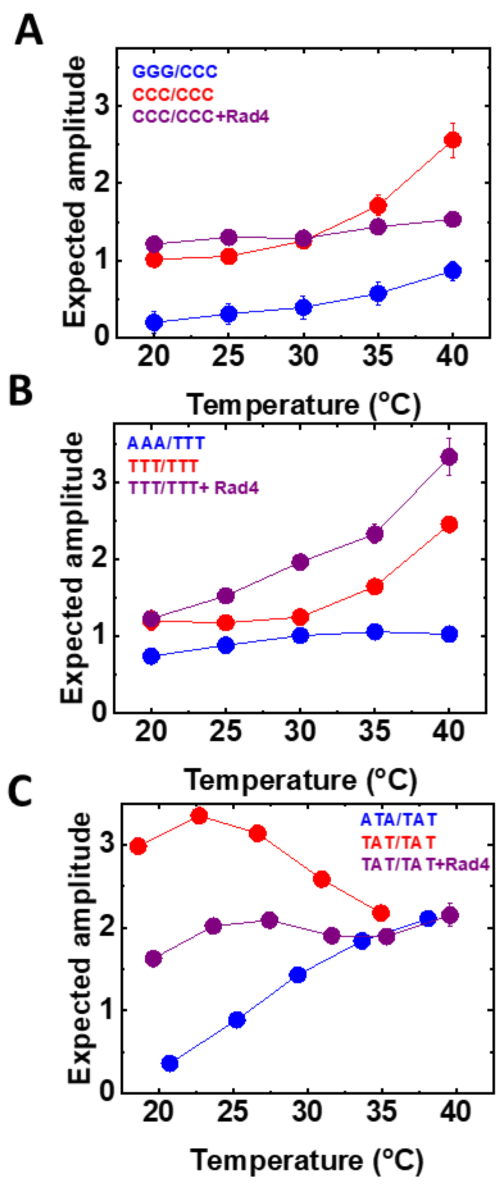


**Figure S11. Comparison of expected amplitudes of conformational relaxation kinetics in response to T-jump perturbation.** The total expected amplitudes for conformational relaxation as computed from equilibrium measurements are plotted as a function of the final temperature after T-jump for (A) the CCC/CCC dataset; (B) the TTT/TTT dataset; and (C) the TAT/TAT dataset. Each panel compares the amplitudes for matched DNA (blue), mismatched DNA (red), and mismatched+Rad4 (purple).
